# GxEsum: a novel approach to estimate the phenotypic variance explained by genome-wide GxE interaction based on GWAS summary statistics for biobank-scale data

**DOI:** 10.1101/2020.05.31.122549

**Authors:** Jisu Shin, S Hong Lee

## Abstract

Genetic variation in response to the environment is fundamental in the biology of complex traits and diseases, i.e. genotype-by-environment interaction (GxE). However, existing methods are computationally demanding and infeasible to handle biobank-scale data. Here we introduce GxEsum, a method for estimating the phenotypic variance explained by genome-wide GxE based on GWAS summary statistics. Through comprehensive simulations and analysis of UK Biobank with 288,837 individuals, we show that GxEsum can handle a large-scale biobank dataset with controlled type I error rates and unbiased GxE estimates, and its computational efficiency can be hundreds of times higher than existing GxE methods.

## Background

The success of the human genome project has led to a paradigm-shift in the complex trait analysis that focuses on the genome-wide association studies (GWAS) [1]. GWAS have been incredibly successful at identifying genome-wide significant single nucleotide polymorphisms (SNPs) that are associated with causal variants underlying complex traits [2, 3]. Moreover, whole-genome approaches, using all common SNPs across the genome, have been useful to dissect the genetic architecture of complex traits, e.g. SNP-based heritability and genetic correlation [4]. However, the analytical modelling used in GWAS and whole-genome approaches usually assumes that there is no genotype-environment interaction (GxE), which can be often violated against the true genetic architecture of complex traits. Indeed, interaction is fundamental in biology and there has been increasing interest in estimating GxE, using genome-wide SNPs [5–7].

Current state-of-the-art whole genome methods for estimating GxE include genotype-covariate interaction genomic restricted maximum likelihood (GREML) and random regression GREML [8]. Recently, a multivariate reaction norm model (RNM) has been introduced [9], which can disentangle GxE from genotype-environmental correlation, providing more reliable GxE estimations. These methods typically employ the GREML approach that requires individual level genotypes and is computationally intensive. Especially when using biobank-scale data, the approach becomes computationally intractable.

To reduce the computational limitation of GREML, linkage disequilibrium score regression (LDSC) was introduced to estimate SNP-based heritability and genetic correlation [10]. LDSC is computationally efficient and requires no individual-level genotypes. Instead, it uses GWAS summary statistics, regressing the association test statistics of SNPs on their LD score. However, existing LDSC methods are limited to additive models only [11–14].

In this study, we propose a novel approach to estimate the phenotypic variance explained by genome-wide GxE based on GWAS summary statistics (GxEsum) for a large-scale biobank dataset, correctly accounting for genotype-environment correlation and scale effects. In simulated and real data analyses, we show that the computational efficiency of the proposed approach is substantially higher than RNM, an existing GREML-based method, while the estimates are reasonably accurate and precise. Because of this computational advantage, GxEsum may be an efficient tool to estimate GxE that can be applied to large-scale data across multiple complex traits.

## Results

### Method Overview

We propose a method to estimate the phenotypic variance explained by the whole-genome GxE, based on GWAS summary statistics, referred to as GxEsum. GxEsum can be a computationally efficient RNM using an extension of the LDSC approach. While the existing LDSC approach is designed to use estimated additive SNP effects in GWAS summary statistics (Supplementary Note 1), GxEsum requires summary statistics of SNP-by-environment interaction effects. For SNP effects modulated by an environment, the expected chi-square statistic 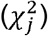 is

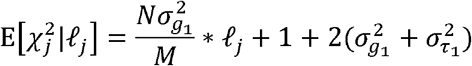

where N is the number of individuals, M is the number of SNPs, 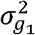 is the variance due to GxE, 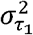 is the variance due to residual heterogeneity or scale effects caused by residual-environment interaction (RxE) and *ℓ_j_* is the LD score at the variant *j* that can be estimated from a reference panel (please see Methods for a full derivation of this equation). The 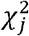 test statistics corresponds to the regression coefficient for the interaction between the *j*th SNP and the environmental covariate (E). The outcome trait is pre-adjusted for confounders and the main effects of E and then a regression model with the main and interaction effects is run by SNP-by-SNP. If chi-square statistics from GWAS are regressed on LD scores, non-genetic interaction effects 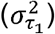 are captured by the intercept, from which GxE 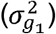 can be disentangled. Consequently, GxE effects estimated by GxEsum are equivalent to that adjusted for RxE when using RNM [9].

To validate the proposed model, i.e. GxEsum, we used various simulations based on real genotype data (see Supplementary Note 2 for a full description of the simulation models). In simulations with and without GxE, we assessed the type l error rates and the accuracy of estimated GxE. We deliberately generated confounding effects such as genotype-environment (G-E) correlation, RxE and residual-environment (R-E) correlation to see if the type l error rate and the accuracy of GxEsum were affected by these confounding factors.

In the real data analysis, we used the UK Biobank data with 288,837 unrelated individuals after stringent quality control. Subsets of the data with various sample sizes were analysed to compare the precision (i.e. power) and the computational efficiency of GxEsum and GREML-based GxE model (i.e. RNM).

Finally, we show how the genetic effects of a complex trait (e.g. BMI, hypertension or type 2 diabetes) are modulated by environment (e.g. neuroticism score, alcohol intake frequency, physical activity or age) by using the proposed method.

### Simulations

For a continuous trait, under the null (no GxE), whether or not there were confounding effects (RxE and G-E and R-E correlations), the type I error rate of GxEsum was not significantly inflated (Table 1). Note that the use of 500 replicates for each simulation scenario can detect a type I error of greater than 0.07 or less than 0.03 as significantly different from 0.05, using the binomial distribution theory [15, 16]. Even with larger confounding effects (Supplementary Table 1), there was no inflation for the type I error rate of GxEsum.

**Table 1.** Type I error rates of GxEsum to detect GxE at a significance threshold of p-value < 0.05.

| Scenarios | Type I Error rate |
| --- | --- |
| Var(GxE<sup>a</sup>) = 0, var(RxE<sup>b</sup>) = 0 | 0.066 |
| Var(GxE) = 0, var(RxE) = 0, G-E correlation<sup>c</sup> = 0.1 | 0.064 |
| Var(GxE) = 0, var(RxE) = 0, R-E correlation<sup>d</sup> = 0.1 | 0.044 |
| Var(GxE) = 0, var(RxE) = 0, G-E correlation=0.1, R-E correlation=0.1 | 0.056 |
| Var(GxE<sup>a</sup>) = 0, var(RxE<sup>b</sup>) = 0.1 | 0.044 |
| Var(GxE) = 0, var(RxE) = 0.1, G-E correlation<sup>c</sup> = 0.1 | 0.034 |
| Var(GxE) = 0, var(RxE) = 0.1, R-E correlation<sup>d</sup> = 0.1 | 0.028 |
| Var(GxE) = 0, var(RxE) = 0.1, G-E correlation=0.1, R-E correlation=0.1 | 0.054 |
| **Average** | **0.049** |

<sup>a</sup>GxE: Genotype-Environment interaction, <sup>b</sup>RxE: Residual-Environment interaction, <sup>c</sup>G-E correlation: Genotype-Environment correlation, <sup>d</sup>R-E correlation: Residual-Environment correlation. We simulated phenotypic data based on a real genotypic dataset (ARIC GWAS) including 7,263 participants with 583,085 SNPs, using various scenarios. The phenotypes were standardised such that the phenotypic mean was 0 and the phenotypic variance was 1. Type I error rate (i.e. false-positive) was estimated from 500 replicates for each scenario.

In simulation with non-zero interactions, estimated GxE (g1) was not remarkably different from the true values whether there were significant G-E and R-E correlations or not (see supplementary Figure 1). It was noted that RxE component was correctly captured by the intercept and not confounded with GxE estimates even when using non-normal environmental variables (Supplementary Note 3 and Supplementary Tables 2 and 3). In the absence of RxE, estimated GxE was also unbiased (Supplementary Figure 2). The estimated GxE seemed robust to different values of G-E and R-E correlations ranging from 0.05 to 0.2, respectively (Supplementary Figures 3 and 4).

On the other hand, estimated main genetic variance (g0) was slightly biased especially when using a large G-E or R-E correlation (Supplementary Figure 4). This is probably because of the fact that the main genetic effects are over-adjusted for the environment due to the large correlations (between the trait and environment) in the model.

We also validated that there was no inflation for the type I error rate when applying GxEsum to binary (disease) traits (Table 2 and Supplementary Table 4), showing that GxEsum appears to be robust to false positives in the scenarios of various confounders. In addition, we estimated the variance component of GxE on the observed scale and transformed to that on the liability scale, using Robertson transformation [17]. As shown in Supplementary Figure 5, the transformed estimates were close to the true simulated values on the liability scale, although the precision of estimates (represented as 95% CI) was shown to be decreased when the population prevalence approached an extreme (e.g. k=0.025). GxE estimates were biased when simulating a large effect size of GxE (e.g. 10% of phenotypic variance explained by GxE) in the case of k=0.025 (Supplementary Figure 6) although they were mostly unbiased in the case of k=0.1 (Supplementary Figure 7). The level of biasedness appeared to be increased when there were RxE effects (Supplementary Figure 6). Finally, caution should be given in interpreting GxE estimates when there are large confounding effects such as substantial G-E and R-E correlations (Supplementary Figure 8). The inflated GxE estimates were probably due to the fact that the phenotypes were over-adjusted for the environment in the model because of the correlation between the main trait and environment (G-E and R-E correlations). This resulted in a reduced phenotypic variance (Supplementary Figure 9), hence an inflated GxE estimates that is the ratio of the estimated GxE variance to the phenotypic variance.

**Table 2.** Type I error rates of GxEsum when using binary disease traits with various population prevalence.

| Scenarios | Population prevalence (k) | Type I error rate |
| --- | --- | --- |
| **Var(GxE) = 0, Var(RxE) = 0** | 0.025 | 0.052 |
|  | 0.05 | 0.042 |
|  | 0.1 | 0.076 |
|  | 0.5 | 0.054 |
| **Var(GxE) = 0, Var(RxE) = 0.1**<br>(on the liability scale) | 0.025 | 0.044 |
|  | 0.05 | 0.036 |
|  | 0.1 | 0.050 |
|  | 0.5 | 0.052 |
| **Average** |  | **0.050** |

We simulated quantitative phenotypic data based on a real genotypic dataset (ARIC GWAS) including 7,263 individuals with 583,085 SNPs. The phenotypes were standardised such that the mean was 0 and variance was 1, for which we applied the liability threshold model to generate affected or unaffected disease status for each individual, using various values for the population prevalence ( $k = 0.025, 0.05, 0.1$ or $0.5$ ). Type I error rate at a significance threshold of $p\text{-value} < 0.05$ was estimated from 500 replicates for each scenario and population prevalence.

Nevertheless, those confounders including RxE interaction and G-E/R-E correlations would not produce false positives whether using continuous quantitative or binary responses as shown above (also see Supplementary Tables 1 – 4). We additionally tested if the type I error rate of GxEsum was controlled when there is collider bias, which is a concern especially when using a self-report study (e.g. UK Biobank data) [18]. In simulations with collider bias, although estimated SNP-heritability was substantially (and unrealistically) underestimated (Supplementary Figures 10 and 11), the type I error rate of GxEsum was well controlled whether using continuous or binary responses (Supplementary Tables 5 and 6).

The estimated variance of the main genetic effects was mostly unbiased when using binary disease traits without G-E/R-E correlations (Supplementary Figure 12). When there were significant G-E and /or R-E correlations, the estimated variance of the main genetic effects appeared to be underestimated especially when there was RxE interaction (Supplementary Figure 13), which confirmed the fact that the main genetic effects are over-adjusted for the environment due to correlations between the trait and environment (Supplementary Figure 9).

### Precision and Computational Efficiency

The precision was assessed by comparing the standard error (SE) of GxEsum and RNM estimates. The SE of GxEsum was obtained from LDSC software (using a jackknife method). The SE of RNM for GxE component can be obtained from the information matrix [19] or from well-established theory [20] (see Supplementary Table 7). Figure 1 shows that the SE of GxEsum was 1.65 times higher than that of RNM when using the same sample size of 50,000. However, when sample size increased for GxEsum up to 288,837, for which RNM estimation is infeasible, the ratio reduced to 0.2. GxEsum can use a larger sample size (e.g. > 1,000,000), for which the ratio is expected to be further decreased, although the largest sample size tested in this study was 288,837 (Supplementary Figure 14).

**Figure 1.**
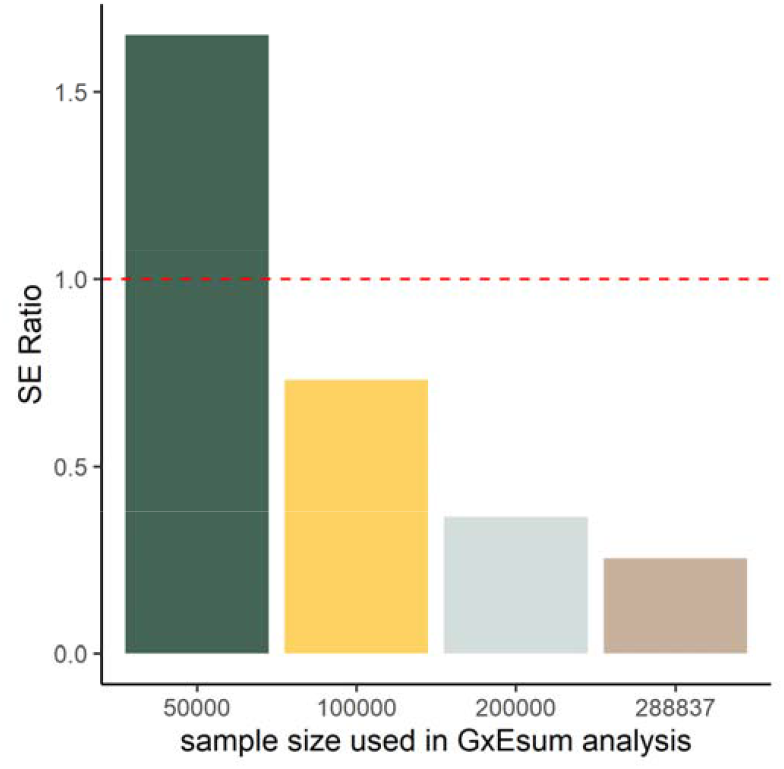
The ratio of standard error (SE) from GxEsum to that from RNM using UK Biobank data. The SEs of GxE variance estimated from GxEsum with various sample sizes ranging from 50,000 to 288,837 were obtained, and they were compared to the SE of GxE variance estimated from RNM with a sample size of 50,000. The dashed horizontal line represents the ratio as 1.

While the precision of GxEsum is competitive with that of RNM, the computational efficiency is dramatically different between two methods (Figure 2 and supplementary Table 8). For example, when using a sample size of 50,000, the computing time for RNM was taken more than a thousand times than GxEsum. Even for GxEsum with a sample size of 288,837, its computational efficiency was still substantially higher than RNM with a sample size of 50,000 (Supplementary Figure 14 and Supplementary Table 7). This justifies that GxEsum is a computationally efficient tool that can be applied to biobank scale data for multiple complex traits and diseases. It is noted that we assumed that preliminary analyses for each method were already done (e.g. GRM for RNM, and LD scores and GWAS for GxEsum) (Supplementary Table 8).

**Figure 2.**
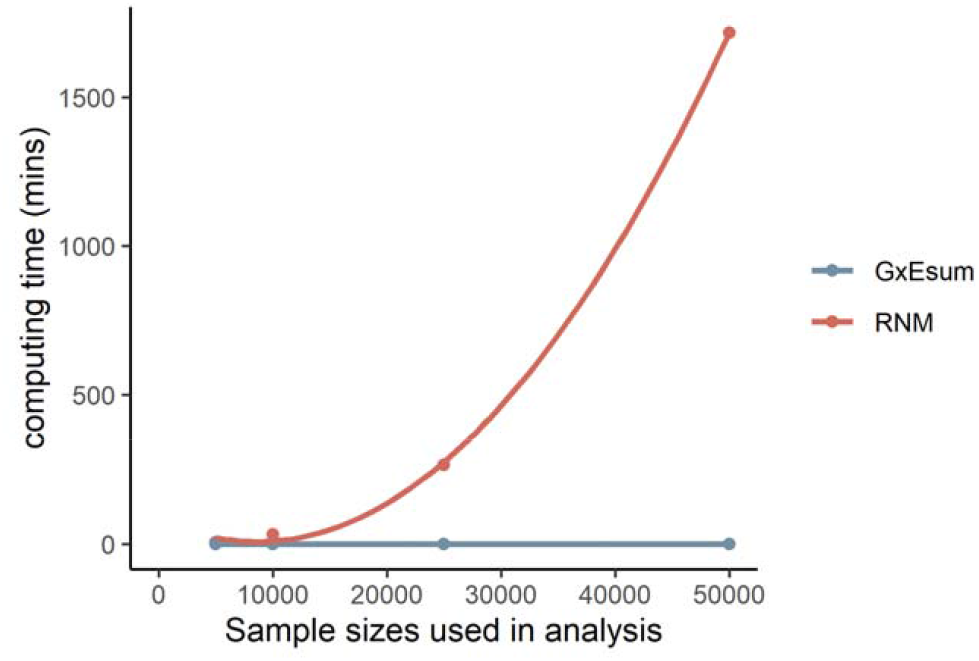
Computing time with various sample sizes used in GxEsum and RNM analyses. As the sample size increases, the computing time of RNM (red) increases exponentially, while that of GxEsum (blue) is almost invariant (less than a minute).

GxEsum uses the Wald test to get a p-value for the null hypothesis, i.e. absence of GxE interaction, using an estimated GxE variance and its standard error. Therefore, the power of the method is closely related to the precision.

### Real data analyses

We applied GxEsum to estimate genetic effects of body mass index (BMI) that were modulated by an environment such as age, alcohol intake frequency, neuroticism scores or physical activity. The significant GxE was observed from the analyses using neuroticism scores. On the other hand, we did not find significant GxE when using age, alcohol intake frequency and physical activity after Bonferroni correction (Bonferroni p-value = 0.05/10 = 0.005 since there were 10 significance tests in this study) (Table 3). The GxEsum approach applied to a binary disease was conducted using hypertension or type 2 diabetes as the main trait, and BMI, waist-hip ratio (WHR). body fat percentage (BFP), or systolic/diastolic blood pressures (BP) as an environmental variable. Table 3 shows that the genetic effects of hypertension and type 2 diabetes were significantly modulated by BMI, but not by other environmental variables. For a comparison, LDSC estimates (i.e. from the null model without GxE) are also shown in Supplementary Table 9.

**Table 3.** Estimates obtained from GxEsum analysis using real data.

| Main trait | Environmental variable | Main additive genetic variance ($\sigma_{g_0}^2$) | GxE interaction variance ($\sigma_{g_1}^2$) | p-value for GxE |
| --- | --- | --- | --- | --- |
| **BMI** | Age | 0.216 (0.007) | 0.004 (0.002) | 1.86E-02 |
|  | NEU<sup>a</sup> | 0.216 (0.007) | 0.007 (0.002) | 1.61E-05 |
|  | PA<sup>b</sup> | 0.218 (0.007) | 0.003 (0.001) | 2.57E-02 |
|  | ALC<sup>c</sup> | 0.216 (0.007) | 0.003 (0.002) | 5.98E-02 |
| **Hypertension** | BMI | 0.152 (0.008) | 0.006 (0.002) | 2.09E-03 |
|  | WHR<sup>d</sup> | 0.154 (0.008) | 0.005 (0.002) | 3.21E-02 |
|  | BFP<sup>e</sup> | 0.151 (0.008) | 0.008 (0.003) | 2.66E-02 |
| **Type 2 Diabetes** | BMI | 0.141 (0.014) | 0.085 (0.022) | 1.58E-04 |
|  | Diastolic BP<sup>f</sup> | 0.198 (0.014) | -0.004 (0.006) | 5.38E-01 |
|  | Systolic BP | 0.204 (0.014) | -0.006 (0.006) | 3.17E-01 |

We used a quantitative trait (BMI) and binary disease traits (hypertension and type 2 diabetes) because BMI is known to be modulated by age/lifestyle such as NEU, ALC, PA [8, 22, 23], and hypertension and type 2 diabetes are known to be caused by obese traits such as BMI and WHR [24, 25]. The p-value is from a Wald test for the estimated GxE variance not being different from zero. The estimates on the observed scale for the binary trait, hypertension, were transformed to those on the liability scale using Robertson transformation[17, 26]. All estimates were from the GxEsum model.
<sup>a</sup>NEU: Neuroticism Score
<sup>b</sup>PA: Physical Activity
<sup>c</sup>ALC: Alcohol intake frequency
<sup>d</sup>WHR: Waist-Hip Ratio
<sup>e</sup>BFP: Body Fat Percentage
<sup>f</sup>BP: Blood pressure

Because not all variables were without missing, we imputed missing phenotypes using the mean value for each variable in the analyses, in order to maximise the sample size. In this real data analysis, there was no remarkable difference in the results whether using phenotypic imputation or not although some variables improved their significance, e.g. NEU (Supplementary Tables 10-12). It is noted that our main phenotypes had a small proportion of missing values, i.e. 0.3%, 7.9% and 0.2% for BMI, hypertension and type 2 diabetes, respectively. If missing rate is substantially high, we recommend to exclude missing values from the analysis, or a better phenotypic imputation method [21] should be used.

## Discussion

In this study, we propose GxEsum, a novel whole-genome GxE method, of which the computational efficiency is a thousand times higher than existing methods. The estimation of GxE using GWAS summary statistics has great flexibility in the application of the method to multiple complex traits and diseases. The proposed method and theory have been explicitly verified using comprehensive simulations that were carried out for both quantitative trait and binary disease. Moreover, we showed that the type I error rate of the proposed method was not inflated by moderate to severe collider bias [18] that caused a substantial underestimation of heritability shown in our simulation (Supplementary Figures 10 and 11).

In the real data analysis, we show that the genetic effects of BMI were significantly modulated by NEU, which agrees with previous studies [9]. It is noted that the significance of GxE was improved because we used a larger sample size, compared with the previous studies. Our result agrees with Robinson et al. (2017) who found no significant GxE evidence for age when analysing BMI using the UK Biobank in which the participants aged 40-69 at the recruitment. However, a dataset with a wider range of ages is desirable, which would increase the power to detect GxE on age. For example, a significant GxE was found in a BMI-age analysis using a dataset including samples aged 18-80 at the recruitment [8]. For hypertension, its causal relationship with BMI has been reported by a number of studies using Mendelian randomization [24, 27]. However, it was not clear if the causal relationship is due to GxE or something else, e.g. unknown non-genetic effects of the disease modulated by BMI status. We show that the causal relationship between hypertension and BMI, and type 2 diabetes and BMI [28] reported in the previous studies [24, 27] may be partly due to genome-wide GxE interaction effects. Interestingly, there is no significant evidence of genome-wide GxE for hypertension-WHR or hypertension-BFP causal relationship that was observed in Mendelian randomization studies [29, 30].

The estimated intercept from GxEsum should be interpreted with caution. We show that estimated intercepts were unbiased from the theoretically predicted values when using the simulation of quantitative traits, as a proof of concept, i.e. the phenotypic variance explained by RxE effects 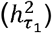 can be obtained as 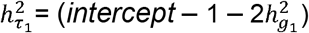 / 2 from eq. (4), or more generally, 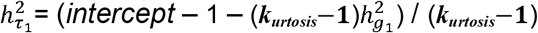 from eq. (5). However, in real data analyses, there may be additional confounding effects such as scale effects, residual heteroscedasticity or/and sample heterogeneity that are often attributed to unknown factors. Moreover, when using binary traits, substantial scale effects can be generated (statistical RxE effects) because only affected and unaffected status are observed and individual differences within affected or unaffected group are ignored. These additional confounding effects and statistical scale effects are captured and estimated as an intercept in GxEsum [10], resulting in unreliable RxE estimates. It is noted that RxE estimation is not the main interest of GxEsum and can be more reliably estimated in RNM that is designed to model both GxE and RxE.

The existing GxE methods require individual-level genotype data which often has a restriction to share, and their computational burden is typically high. Moreover, it is not clear how they perform when the representativeness of the samples is limited, e.g. selection bias due to a collider in the UK Biobank samples. On the contrary, the proposed approach, GxEsum, is computationally efficient and can detect GxE interaction correctly for both quantitative and binary disease traits even when there is moderate to server collider bias. If GWAS summary statistics of estimated main additive and interaction effects can be made publicly available, a meta-analysis across multiple cohorts can be possible for an ever-large GxE study (like the context of LDSC SNP-heritability meta-analysis). There are some issues that the measure of environmental variable may not be standardised across study cohorts, and the environmental variable may be even unavailable in some cohorts. However, these issues can be remedied when the information of exposome that is the standardised measure of all exposures for individuals, complemented to the genome, is available.

There is a GxE method that can use GWAS summary statistics, i.e. VarExp, which is recently published. While VarExp benefits computationally from using GWAS summary statistics, it needs to invert the correlation matrix between SNPs, which prevents from using a large number of SNPs [31]. Furthermore, the theoretical frameworks of GxEsum and VarExp are fundamentally different in that the latter does not account for confounding effects such as scale effects, residual heterogeneity or RxE that can be captured by the estimated intercept of GxEsum. Finally, the performance of VarExp has been verified with a limited magnitude of interaction effects up to 1.5% and 0.25% of the phenotypic variance for quantitative and binary traits, respectively [31].

Like RNM, GxEsum can fit environmental exposures such that the genetic effects of a trait can be modelled as a nonlinear function of a continuous environmental gradient. The potential modifier of the genetic effects is not limited to environmental exposures but can be extended to novel variables from multi-omics data such as gene-expression, protein expression and methylation data [32, 33]. Polygenic risk scores [34, 35] can also be considered as an environmental variable in the model. This novel approach may allow dissecting a latent biological architecture of a complex trait in a future application of GxEsum.

In the analysis of binary disease traits, estimates on the liability scale, transformed from those on the observed scale using Robertson transformation, should be interpreted with caution. Biased estimates on the liability scale are likely due to the violation of the normality assumption that is essentially required for the Robertson transformation, i.e. large interaction effects can cause a non-normal phenotypic distribution. It is also known that if the transformation involves substantial non-additive effects, it can give biased estimates on the liability scale [17, 26, 36]. However, when non-additive effects are small, the transformation can give reasonably accurate estimates on the liability scale, which is also evidenced by our simulations with small interaction effects. As shown in the real data analysis, the magnitude of genome-wide GxE is not large (< 10% of the phenotypic variance), showing that the bias of transformation due to the assumption violation may not be substantial in general. Nevertheless, it is required to develop a better transformation method for large interaction effects in a further study, e.g. when using multiple environmental variables simultaneously, the interaction effects are aggregated and can be substantially large.

There are a number of limitations in our study. First, like RNM, GxEsum does not determine the causal direction between variables, which can be provided from previous studies or other epidemiologic methods, e.g. Mendelian randomisation, as prior information. Second, we only modelled the first order of random regression coefficients with a single environmental variable, and there may be significant additional effects when modelling a higher-order interaction or multiple environmental variables. It is possible to extend GxEsum model to fit additional quadratic and polynomial terms or multiple environmental variables simultaneously. However, assessing the performance of these advanced models is a formidable task, requiring a further study. Third, the estimation for the main genetic effects can be biased when there are large G-E and/or R-E correlations. Because of such correlations, the main genetic effects are over-adjusted when the phenotypes of the main trait are adjusted for the environmental variable in the model. Therefore, a careful interpretation of the estimated main genetic effects is required when using GxEsum. Fourth, we did not investigate the performance of GxEsum for ascertained case-control studies in which cases are over-sampled. A further study is required to extend the method to non-random case-control samples so that it can be applied to consortium data with multiple case-control studies. Lastly, when using the same sample size, the precision of GxEsum is not better than GREML-based GxE methods, implying that the former is only useful when using a large sample size that the latter cannot handle.

## Conclusions

Despite these caveats, GxEsum can be a useful tool to estimate whole-genome GxE as it can achieve a higher precision (i.e. power) from a larger sample size, compared to existing GxE methods. Especially when the scale of available resources increases, GxEsum may be a unique method that can be applied to large-scale data across multiple complex traits and diseases in the context of GxE.

## Methods

### GxEsum

Following Ni et al. (2019), RNM can be written as

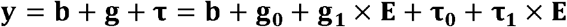

where y is *N* vector of phenotypic observations, **b** is a vector of fixed effects, **g** is *N* individual genetic effects, which can be decomposed into the first and second order of genetic random regression coefficients, **g**_0_ and **g_1_, τ** is residual effects, decomposed into the first and second order of residual random regression coefficients, **τ_0_** and **τ_1_**, and E is an *N* vector of environmental variable. Note that **E** can be also any covariate variable (e.g. smoking, alcohol intake frequency).

Assuming that the phenotypes (**y**) are pre-adjusted for the main genetic effects (**g_0_**), environmental or covariate variable (**E**) and other fixed effects (**b**), the model can be rewritten as

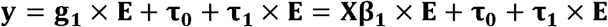

where X is an *N* x *M* standardised genotype matrix for *M* SNPs, **β_1_** is an *M* vector of SNP interaction effects modulated by the environment (i.e. GxE SNP effects). It is loted that **τ_0_** is residual effects that are consistent across environment whereas *τ_1_* captures heterogeneous residual effects across environment (i.e. RxE).

Following Bulik-Sullivan et al. (2015), assuming 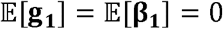, the expected chisquare statistics of variant *j* for the GxE is

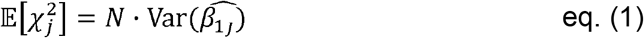

Using the law of total variance, 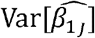 can be obtained as

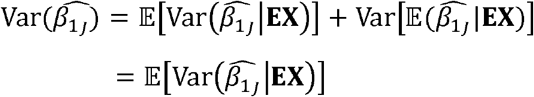

where **EX** is an *N* x *M* matrix with each column having the Hadamard product between **E** and **X**_*j*_ (standardised genotypes at the *j* th SNP) and the conditional expectation of 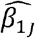 is 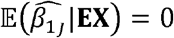.

Noting that the least-square estimate of 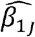 can be obtained as 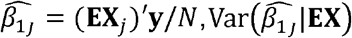 can be rearranged as

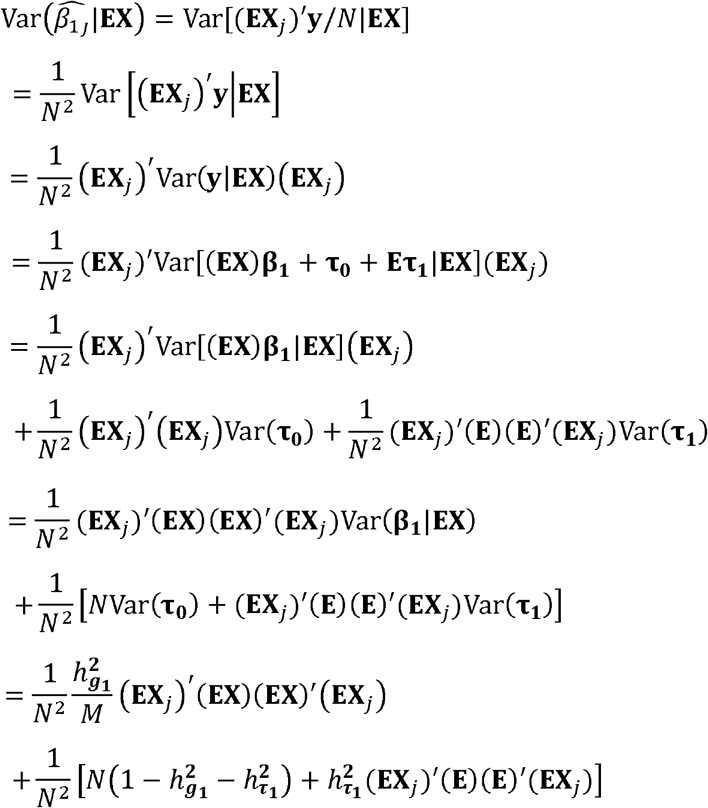

where 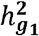 and 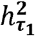 is the proportion of phenotypic variance explained by GxE and RxE, respectively.

Therefore, 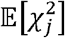 in eq. (1) can be written as

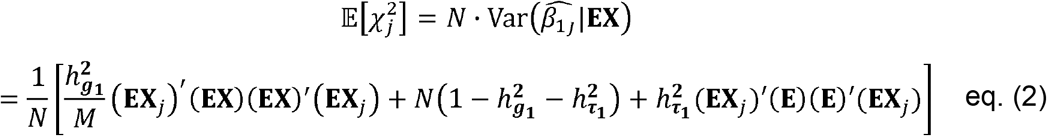

According to Bulik-Sullivan et al. (2015), the products of the standardised genotypes at variant *j* and other variants can be expressed as a function of LD scores, i.e.

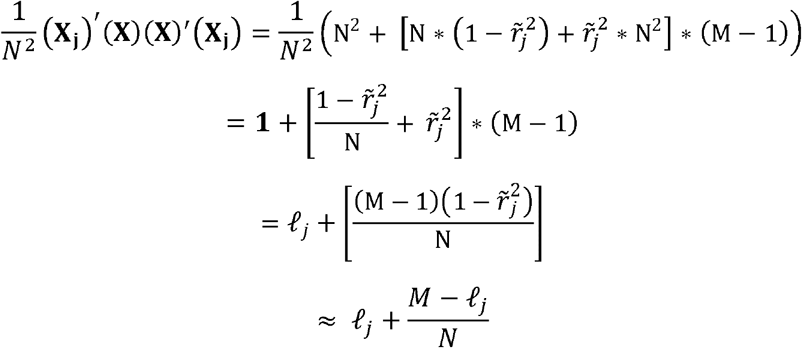

where 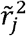 defined as the expected sample correlation between genotypes at the *j*th variant and the other (*M*-1) variants, and 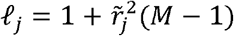 is the LD scores of the *j*th SNP.

According to the central moment theory of standard normal distribution of three independent random variables (**X_1_, X_2_** and **E**), each with an N vector, useful equations are

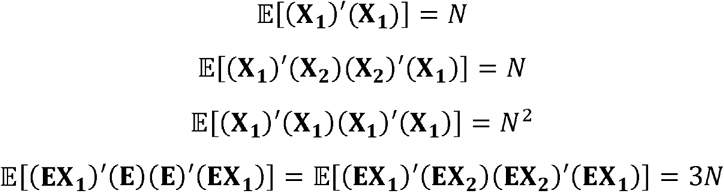

and

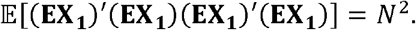

Therefore, assuming that **E** and **X_j_** have negligible correlation for a polygenic trait (i.e. a tiny proportion of the phenotypic variance of E can be explained by a single SNP, X_*j*_), the term (**EX**_*j*_)’(**EX**)(**EX**)’(**EX**_*j*_) can be expressed as a function of LD scores as

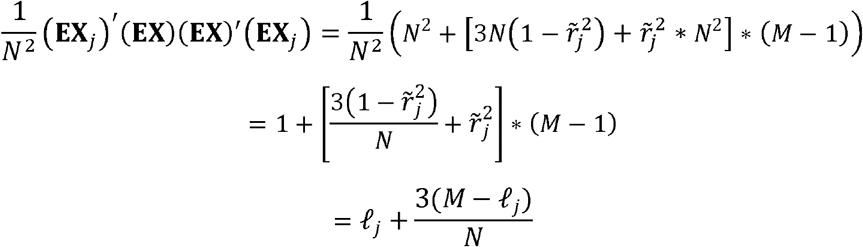

Thus, a part in eq. (2) can be rearranged as

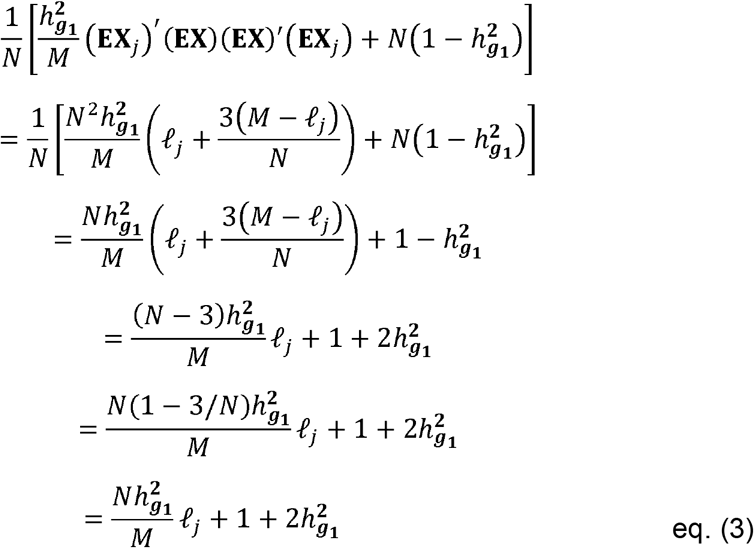

The term, 1 – 3/*N*, in eq. (3) can be approximated as 1 in the analysis using biobank scale data, which contains over 10^5^ samples.

The remaining part in eq. (2) can be rearranged as

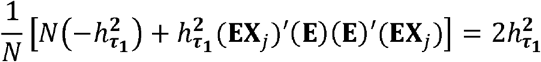

where 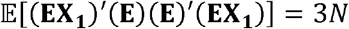 according to the central moment theory of standard normal distribution (see above), assuming that E and each column of **X** have a negligible correlation, which satisfies if E is an environmental variable or a polygenic trait.

Therefore,

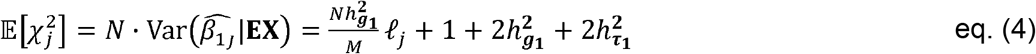

where 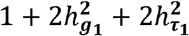 can be obtained as the intercept of the outcome by fitting to the proposed model (GxEsum). It is noted Eq. (4) is valid when **X**_*j*_ is not strictly normally distributed, i.e. the centred and standardised genotypes of jth SNP, which is already shown in Bulik-Sullivan et al. [10]. When the environmental variable (**E**) is non-normal, the general form of the fourth central moment term can be expressed as 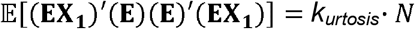 where *k_urtosis_* is the kurtosis of **E**. And, only the intercept part of the Eq. (4) is slightly modified as

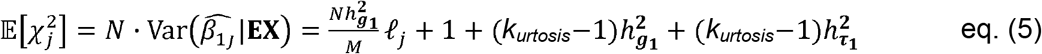

Eq. (4) and (5) are verified using simulations (see Supplementary Note 3 and Supplementary Table 2).

To validate the proposed model in general, we used comprehensive phenotypic simulations that were based on real genotype data (see Supplementary Note 4).

### Real data

UK Biobank data were used, which contains 0.5 million individuals aged between 40-69 years. The data consists of health-related information for each participant who was recruited in 2006-2010, and their imputed genomic data (~92 million SNPs) has been distributed through European Genome-phenome Archive. A stringent quality control process for individuals was set as followings: 1) who were reported as non-white British, 2) who were having mismatched gender between the reported and the inferred by the genotypic data, 3) who were having missing rate over 0.05, 4) who were having putative sex chromosome aneuploidy. In addition, only HapMap3 SNPs were used which were passed from the stringent quality controls for SNPs. The filter for SNPs is set as followings: 1) which were having INFO score less than 0.6, 2) which were having a MAF less than 1%, 3) which were having Hardy-Weinberg Equilibrium (HWE) P-value less than 1E-4, 4) one of which from the duplicated SNPs. From those passing the tough procedures, we additionally excluded one of pair of samples who were having the genomic relationship higher than 0.05. After quality control, 288,837 individuals and 1,133,273 SNPs were remained. We estimated LD scores using the genotypic data of UK Biobank after these quality control processes.

Among trait phenotypes available in the UK biobank, we arbitrarily selected BMI (a quantitative trait), hypertension and type 2 diabetes (binary disease traits) and tested if the genetic effects of the complex traits were significantly modulated by an environmental variable, i.e. NEU, ALC, PA or age (for testing BMI), BMI, WHR or BFP (for hypertension), and BMI, diastolic BP or systolic BP (for type 2 diabetes). The number cases for hypertension and type 2 diabetes was 134,499 (population prevalence is 0.51) and 11,694 (population prevalence is 0.04), respectively. The phenotypes of the main trait were adjusted for potential confounders such as age, gender, year of birth, assessment centre, Townsend Deprivation Index, genetic batch, household income, educational qualification, the first 10 principal components, and the environmental variable. For any phenotypic missing value for each variable, we used the mean of the phenotypes of the variable, i.e. phenotypic imputation with the mean. A better phenotypic imputation method [21] can be used, which is likely to improve the significance of GxE. Further details of the variables used on this study are in Supplementary Note 4.

In GWAS, we used a linear model for quantitative traits as well as for binary responses. The use of a linear model applied to binary responses is because it has been reported that a logistic regression may generate biased estimates in some instances [37] and our simulations (Supplementary Note 2) were based on a probit model (i.e. a linear transformation of the inverse standard normal distribution) that can be well approximated by a linear model [38].

## Abbreviations

GxE: genotype-by-environment interaction
RNM: reaction norm model
GWAS: genome-wide association studies
SNPs: single nucleotide polymorphisms
GREML: genomic restricted maximum likelihood
LDSC: linkage disequilibrium score regression
RxE: residual-environment interaction
G-E correlation: genotype-environment correlation
R-E correlation: residual-environment correlation
SE: standard error
BMI: body mass index
WHR: waist-hip ratio
BFP: body fat percentage
NEU: neuroticism score
PA: physical activity
ALC: alcohol intake frequency

## Acknowledgements

The HPC resources were provided by the Australian Government through Gadi under the National Computational Merit Allocation Scheme (NCMAS) and by the university of South Australia through Tango 2.0. We would like to thank staff and participants of the ARIC study and the UK Biobank for their valuable contributions. The Atherosclerosis Risk in Communities Study is carried out as a collaborative study supported by National Heart, Lung, and Blood Institute contracts (HHSN268201100005C, HHSN268201100006C, HHSN268201100007C, HHSN268201100008C, HHSN268201100009C, HHSN268201100010C, HHSN268201100011C, and HHSN268201100012C). We thank the staff and participants of the ARIC study for their important contributions. Funding for GENEVA was provided by National Human Genome Research Institute grant U01HG004402 (E. Boerwinkle).

## Authors’ contribution

S.H.L. conceived the idea and directed the study. S.H.L. and J.S. derived and verified the theory. J.S performed the analyses. J.S. and S.H.L. drafted the manuscript.

## Availability of Data and Materials

### Data

The UK Biobank data are accessed via https://www.ukbiobank.ac.uk/

The ARIC study data are accessed via dbGaP (https://dbgap.ncbi.nlm.nih.gov) and its accession code is phs000280.v7.p1.

### Software

GxEsum model is implemented in the script that are publicly available at https://github.com/honglee0707/GxEsum, and the demonstration of GxEsum software is described in Supplementary Note 5. The version of source code used in the manuscript is deposited with DOI: 10.5281/zenodo.4659681 at https://zenodo.org/record/4659681#.YGkZXc9xeUk.

LDSC can be download from https://github.com/bulik/ldsc

PLINK version 1.9 can be download from https://www.cog-genomics.org/plink/1.9/

MTG2 version 2.15 can be download from https://sites.google.com/site/honglee0707/mtg2

## Ethics approval and consent to participant

The current study was approved by the University of South Australia Human Research Ethics Committee. The ARIC Study was approved by the institutional review boards of all participating institutions, including the University of Minnesota, Johns Hopkins University, University of North Carolina, University of Mississippi Medical Centre, and Wake Forest University. The UK Biobank was approved by the North West Multi-centre Research Ethics Committee (11/NW/0382). the reference number approved by the UK Biobank is 14575. All UK Biobank and ARIC Study participants gave written informed consent.

