## Supplementary information for "GxEsum: a novel approach to estimate the phenotypic variance explained by genome-wide GxE interaction based on GWAS summary statistics for biobank-scale data"

**Supplementary Note 1:** LDSC
**Supplementary Note 2:** Simulated data
**Supplementary Note 3:** Theory Verification
**Supplementary Note 4:** Details of data obtained from UK Biobank
**Supplementary Note 5:** Demonstration of GxEsum software

**Supplementary Figure 1**: Estimated variance components of main genetic and GxE effects using GxE sum in the presence of GxE and RxE (i.e. Full scenario)
**Supplementary Figure 2**: Estimated variance components of main genetic and GxE effects using GxEsum in the presence of GxE without RxE (i.e. GxE only scenario).
**Supplementary Figure 3**: Estimated variance components of main genetic and GxE effects using GxEsum when the possible correlations are smaller effects in the presence of both interactions (i.e. full scenario).
**Supplementary Figure 4**: Estimated variance components of main genetic and GxE effects using GxEsum when the possible correlations are larger effects in the presence of both interactions (i.e. full scenario).
**Supplementary Figure 5**: Estimated variance component of GxE when binary diseases traits were used.
**Supplementary Figure 6:** The level of biasness increases when the proportion of phenotypic variance explained by GxE increases in the case of population prevalence k=0.025
**Supplementary Figure 7:** The level of biasness increases when the proportion of phenotypic variance explained by GxE increases in the case of population prevalence k=0.1
**Supplementary Figure 8:** Estimated variance component for GxE when binary disease traits (k=0.1) with which environmental variable is significantly correlated (i.e. G-E and/or R-E correlations).
**Supplementary Figure 9**: The decreasing rate of phenotypic variances after adjusting for the environment.
**Supplementary Figure 10:** Estimated variance of the main genetic effects using quantitative trait when the collider bias was considered.
**Supplementary Figure 11:** Estimated variance of main genetic effects using binary trait (k=0.1) when the collider bias was considered.
**Supplementary Figure 12:** Estimated variance component of the main genetic effects (g0) when binary disease traits were used.
**Supplementary Figure 13**: Estimated variance component of the main genetic effects (g0) when further potential confounders were considered in the case of population prevalence k=0.1.
**Supplementary Figure 14:** The ratio of SE from GxEsum to that from RNM using UK Biobank.

**Supplementary Table 1**: The type I error rates of GxEsum when confounding effects are large.
**Supplementary Table 2**: Theory verification using the obtained intercept values
**Supplementary Table 3:** Type I error rates of GxEsum when using binary environmental variable (k=0.1)
**Supplementary Table 4**: Type I error rates when using binary disease traits (k=0.1) with various confounders.
**Supplementary Table 5:** Type I error rates of GxEsum when the collider bias was introduced in simulated quantitative traits.

**Supplementary Table 6:** Type I error rates of GxEsum in the presence of collider bias when using binary diseases trait (k=0.1).
**Supplementary Table 7**: Comparison of the standard error (SE) of estimated GxE variance, obtained from the GCTA-GREML power calculator and from the information matrix in the RNM.
**Supplementary Table 8**: Comparison of computing time between RNM and GxEsum approaches in the GxE estimation.
**Supplementary Table 9**: Estimates obtained from LDSC and GxEsum analyses using real data
**Supplementary Table 10**: Obtained p-values for GxE in BMI with 4 covariates with and without phenotypic imputation
**Supplementary Table 11**: Obtained p-values for GxE in hypertension with 3 covariates with and without phenotypic imputation
**Supplementary Table 12**: Obtained p-values for GxE Type 2 diabetes with 3 covariates with and without phenotypic imputation

**References**

### Supplementary Note

#### 1. LDSC

Bulik-Sullivan et al. (2015) proposed a model to estimate a SNP-based heritability based on GWAS summary statistics of estimated additive genetic effects. The model can be written as

$$\mathbf{y} = \mathbf{X}\boldsymbol{\beta} + \mathbf{e}$$

where $\mathbf{y}$ is a vector of *N* phenotypic observations, $\mathbf{X}$ is an $N$ x $M$ standardised genotype matrix for *N* individuals and $M$ SNPs, $\boldsymbol{\beta}$ is a vector of *M* SNP effects and $\mathbf{e}$ is residual. The estimated effect size of the $j$th SNP variant (i.e. $\hat{\beta}_j$) can be expressed as

$$\hat{\beta}_j = \mathbf{X}_j^{\mathrm{T}}\mathbf{y}/N$$

where $\mathbf{X}_j^{\mathrm{T}}$ is the transpose of the jth column vector in the genotype matrix, i.e. genotypic information for $N$ individuals at the jth SNP variant. In Bulik-Sullivan et al. (2015), the expected chi-squared statistics of variant $j$ can be derived as a function of LD scores as

$$\begin{aligned}\mathrm{E}[\chi_j^2] &= \mathrm{N} * \mathrm{var}[\widehat{\beta_j}] \\ &= \frac{N * h_\beta^2}{M} * \ell_j + 1\end{aligned}$$

where $h_\beta^2$ indicates the proportion of phenotypic variance explained by the additive genetic effects (i.e. SNP-based heritability) that can be estimated by regressing the chi-squared statistics on LD score ($\ell_j$) that can be estimated from a reference panel or in-sample.

#### 2. Simulated data

##### Quantitative traits

The various phenotype simulations were conducted based on the real genotypic data from the Atherosclerosis Risk in Communities (ARIC) study, which contains 583,085 SNPs from 7,263 unrelated individuals (i.e. any pairwise relatedness < 0.05) that went through the standard quality control. In the quality control process, SNPs were excluded if their minor allele frequency (MAF) were less than 0.01, call rates were less than 0.95 and Hardy-Weinberg equilibrium p-values were less than 0.001, and individuals were excluded if their genotype call rates were less than 0.95.

The simulation model can be written as

$$\mathbf{y} = \mathbf{g_0} + \mathbf{g_1} \times \mathbf{E} + \boldsymbol{\tau_0} + \boldsymbol{\tau_1} \times \mathbf{E}$$
$$\mathbf{E} = \boldsymbol{\gamma} + \boldsymbol{\varepsilon}$$

where $\mathbf{y}$ is *N* vector of phenotypic observations, $\mathbf{g_0}$ and $\mathbf{g_1}$ are the first and second order of genetic random regression coefficients, and $\boldsymbol{\tau_0}$ and $\boldsymbol{\tau_1}$ are the first and second order of random residual random regression coefficients. $\mathbf{E}$ is an *N* vector of environmental variables, and this can be decomposed into $\boldsymbol{\gamma}$ and $\boldsymbol{\varepsilon}$ which are the random genetic and residual effects of environmental variable $\mathbf{E}$. The variance and covariance structure of genetic effects can be represented as

$$\mathrm{cov}(\mathbf{g_0},\mathbf{g_1},\boldsymbol{\gamma}) = \begin{bmatrix} \mathrm{var}(\mathbf{g_0}) & \mathrm{cov}(\mathbf{g_0},\mathbf{g_1}) & \mathrm{cov}(\mathbf{g_0},\boldsymbol{\gamma}) \\ \mathrm{cov}(\mathbf{g_0},\mathbf{g_1}) & \mathrm{var}(\mathbf{g_1}) & \mathrm{cov}(\mathbf{g_1},\boldsymbol{\gamma}) \\ \mathrm{cov}(\mathbf{g_0},\boldsymbol{\gamma}) & \mathrm{cov}(\mathbf{g_1},\boldsymbol{\gamma}) & \mathrm{var}(\boldsymbol{\gamma}) \end{bmatrix} = \begin{bmatrix} \sigma^2_{\mathbf{g_0}} & \sigma_{\mathbf{g_0},\mathbf{g_1}} & \sigma_{\mathbf{g_0},\boldsymbol{\gamma}} \\ \sigma_{\mathbf{g_0},\mathbf{g_1}} & \sigma^2_{\mathbf{g_1}} & \sigma_{\mathbf{g_1},\boldsymbol{\gamma}} \\ \sigma_{\mathbf{g_0},\boldsymbol{\gamma}} & \sigma_{\mathbf{g_1},\boldsymbol{\gamma}} & \sigma^2_{\boldsymbol{\gamma}} \end{bmatrix}$$

where $\sigma^2_{\mathbf{g_0}}$ and $\sigma^2_{\mathbf{g_1}}$ are the variances of the main genetic and GxE effects of the main trait, and $\sigma^2_{\boldsymbol{\gamma}}$ is the genetic variance of the environmental variable. The covariance term, $\sigma_{\mathbf{g_0},\mathbf{g_1}}$, indicates the genetic covariance between the main genetic and GxE effects of the main trait, $\sigma_{\mathbf{g_0},\boldsymbol{\gamma}}$ is the covariance of the main genetic effects between the main trait and environmental variable, and $\sigma_{\mathbf{g_1},\boldsymbol{\gamma}}$ is the covariance between the GxE effect of the main trait and the main genetic effects of the environmental variable.

The variance and covariance structure of residual effects can be represented as

$$\mathrm{cov}(\boldsymbol{\tau_0},\boldsymbol{\tau_1},\boldsymbol{\varepsilon}) = \begin{bmatrix} \mathrm{var}(\boldsymbol{\tau_0}) & \mathrm{cov}(\boldsymbol{\tau_0},\boldsymbol{\tau_1}) & \mathrm{cov}(\boldsymbol{\tau_0},\boldsymbol{\varepsilon}) \\ \mathrm{cov}(\boldsymbol{\tau_0},\boldsymbol{\tau_1}) & \mathrm{var}(\boldsymbol{\tau_1}) & \mathrm{cov}(\boldsymbol{\tau_1},\boldsymbol{\varepsilon}) \\ \mathrm{cov}(\boldsymbol{\tau_0},\boldsymbol{\varepsilon}) & \mathrm{cov}(\boldsymbol{\tau_1},\boldsymbol{\varepsilon}) & \mathrm{var}(\boldsymbol{\varepsilon}) \end{bmatrix} = \begin{bmatrix} \sigma^2_{\boldsymbol{\tau_0}} & \sigma_{\boldsymbol{\tau_0},\boldsymbol{\tau_1}} & \sigma_{\boldsymbol{\tau_0},\boldsymbol{\varepsilon}} \\ \sigma_{\boldsymbol{\tau_0},\boldsymbol{\tau_1}} & \sigma^2_{\boldsymbol{\tau_1}} & \sigma_{\boldsymbol{\tau_1},\boldsymbol{\varepsilon}} \\ \sigma_{\boldsymbol{\tau_0},\boldsymbol{\varepsilon}} & \sigma_{\boldsymbol{\tau_1},\boldsymbol{\varepsilon}} & \sigma^2_{\boldsymbol{\varepsilon}} \end{bmatrix}$$

where $\sigma^2_{\boldsymbol{\tau_0}}$ and $\sigma^2_{\boldsymbol{\tau_1}}$ are the variances of the residual and RxE effects of the main trait, $\sigma^2_{\boldsymbol{\varepsilon}}$ is the residual variance of the environmental variable. The covariance term, $\sigma_{\boldsymbol{\tau_0},\boldsymbol{\tau_1}}$, indicates the covariance between the residual and RxE effects of the main trait, $\sigma_{\boldsymbol{\tau_0},\boldsymbol{\varepsilon}}$ is the residual covariance between the main trait and environmental variable, and $\sigma_{\boldsymbol{\tau_1},\boldsymbol{\varepsilon}}$ is the covariance between the RxE effect of the main trait and the residual effects of the environmental variable.

Based on the simulation model, we tested various scenarios altering the values of simulation parameters, e.g. situations in the absence and presence of GxE with and without the confounders such as RxE and G-E/R-E correlations.

In the null simulation (in the absence of GxE and RxE), the variance and covariance matrix of genetic effects was set as

$$\mathrm{cov}(\mathbf{g_0},\mathbf{g_1},\boldsymbol{\gamma}) = \begin{bmatrix} \sigma^2_{\mathbf{g_0}} & \sigma_{\mathbf{g_0},\mathbf{g_1}} & \sigma_{\mathbf{g_0},\boldsymbol{\gamma}} \\ \sigma_{\mathbf{g_0},\mathbf{g_1}} & \sigma^2_{\mathbf{g_1}} & \sigma_{\mathbf{g_1},\boldsymbol{\gamma}} \\ \sigma_{\mathbf{g_0},\boldsymbol{\gamma}} & \sigma_{\mathbf{g_1},\boldsymbol{\gamma}} & \sigma^2_{\boldsymbol{\gamma}} \end{bmatrix} = \begin{bmatrix} 0.5 & 0 & \sigma_{\mathbf{g_0},\boldsymbol{\gamma}} \\ 0 & 0 & 0 \\ \sigma_{\mathbf{g_0},\boldsymbol{\gamma}} & 0 & 0.5 \end{bmatrix}$$

where $\sigma_{\mathbf{g_0},\boldsymbol{\gamma}}$ (the genetic correlation between the main trait and environment) varied from 0 to 0.25 according to the tested scenario. And, the variance and covariance matrix of residual effects was set as

$$\mathrm{cov}(\boldsymbol{\tau_0},\boldsymbol{\tau_1},\boldsymbol{\varepsilon}) = \begin{bmatrix} \sigma^2_{\boldsymbol{\tau_0}} & \sigma_{\boldsymbol{\tau_0},\boldsymbol{\tau_1}} & \sigma_{\boldsymbol{\tau_0},\boldsymbol{\varepsilon}} \\ \sigma_{\boldsymbol{\tau_0},\boldsymbol{\tau_1}} & \sigma^2_{\boldsymbol{\tau_1}} & \sigma_{\boldsymbol{\tau_1},\boldsymbol{\varepsilon}} \\ \sigma_{\boldsymbol{\tau_0},\boldsymbol{\varepsilon}} & \sigma_{\boldsymbol{\tau_1},\boldsymbol{\varepsilon}} & \sigma^2_{\boldsymbol{\varepsilon}} \end{bmatrix} = \begin{bmatrix} 0.5 & 0 & \sigma_{\boldsymbol{\tau_0},\boldsymbol{\varepsilon}} \\ 0 & 0 & 0 \\ \sigma_{\boldsymbol{\tau_0},\boldsymbol{\varepsilon}} & 0 & 0.5 \end{bmatrix}$$

where $\sigma_{\boldsymbol{\tau_0},\boldsymbol{\varepsilon}}$ (residual correlation between the main trait and environment) varied from 0 to 0.25.

In the absence of GxE with significant RxE effects (denoted as RxE only scenario), the variance and covariance matrices of genetic and residual effects were set as

$$\mathrm{cov}(\mathbf{g_0}, \mathbf{g_1}, \boldsymbol{\gamma}) = \begin{bmatrix} \sigma^2_{\mathbf{g_0}} & \sigma_{\mathbf{g_0},\mathbf{g_1}} & \sigma_{\mathbf{g_0},\boldsymbol{\gamma}} \\ \sigma_{\mathbf{g_0},\mathbf{g_1}} & \sigma^2_{\mathbf{g_1}} & \sigma_{\mathbf{g_1},\boldsymbol{\gamma}} \\ \sigma_{\mathbf{g_0},\boldsymbol{\gamma}} & \sigma_{\mathbf{g_1},\boldsymbol{\gamma}} & \sigma^2_{\boldsymbol{\gamma}} \end{bmatrix} = \begin{bmatrix} (1-\sigma^2_{\mathbf{g_1}}-\sigma^2_{\boldsymbol{\tau}_1})/2 & 0 & \sigma_{\mathbf{g_0},\boldsymbol{\gamma}} \\ 0 & 0 & 0 \\ \sigma_{\mathbf{g_0},\boldsymbol{\gamma}} & 0 & 0.5 \end{bmatrix}$$

and

$$\mathrm{cov}(\boldsymbol{\tau_0}, \boldsymbol{\tau_1}, \boldsymbol{\varepsilon}) = \begin{bmatrix} \sigma^2_{\boldsymbol{\tau_0}} & \sigma_{\boldsymbol{\tau_0},\boldsymbol{\tau_1}} & \sigma_{\boldsymbol{\tau_0},\boldsymbol{\varepsilon}} \\ \sigma_{\boldsymbol{\tau_0},\boldsymbol{\tau_1}} & \sigma^2_{\boldsymbol{\tau_1}} & \sigma_{\boldsymbol{\tau_1},\boldsymbol{\varepsilon}} \\ \sigma_{\boldsymbol{\tau_0},\boldsymbol{\varepsilon}} & \sigma_{\boldsymbol{\tau_1},\boldsymbol{\varepsilon}} & \sigma^2_{\boldsymbol{\varepsilon}} \end{bmatrix} = \begin{bmatrix} (1-\sigma^2_{\mathbf{g_1}}-\sigma^2_{\boldsymbol{\tau}_1})/2 & 0 & \sigma_{\boldsymbol{\tau_0},\boldsymbol{\varepsilon}} \\ 0 & \sigma^2_{\boldsymbol{\tau_1}} & 0 \\ \sigma_{\boldsymbol{\tau_0},\boldsymbol{\varepsilon}} & 0 & 0.5 \end{bmatrix}$$

where $\sigma^2_{\mathbf{g_0}}$ and $\sigma^2_{\boldsymbol{\tau_0}}$ were $(1-\sigma^2_{\mathbf{g_1}}-\sigma^2_{\boldsymbol{\tau_1}})/2$, it can be represented as $(1-\sigma^2_{\boldsymbol{\tau_1}})/2$ in the absence of GxE effect. $\sigma^2_{\boldsymbol{\tau_1}}$ was 0.1 for the rest of the scenarios except for those with large effects size, therefore, $\sigma^2_{\mathbf{g_0}}$ and $\sigma^2_{\boldsymbol{\tau_0}}$ are 0.45, respectively, and $\sigma_{\mathbf{g_0},\boldsymbol{\gamma}}$ and $\sigma_{\boldsymbol{\tau_0},\boldsymbol{\varepsilon}}$ were considered from 0 to 0.2 in accordance with the simulation scenario. In simulations using large confounding effects, $\sigma^2_{\boldsymbol{\tau_1}}$ was 0.34, therefore, the variances explained by the main genetic effects and residual effects (i.e., $\sigma^2_{\mathbf{g_0}}$ and $\sigma^2_{\boldsymbol{\tau_0}}$) are 0.33, respectively, and $\sigma_{\mathbf{g_0},\boldsymbol{\gamma}}$ and $\sigma_{\boldsymbol{\tau_0},\boldsymbol{\varepsilon}}$ varied from 0 to 0.2.

In the presence of GxE without RxE (denoted as GxE only scenario), the variance and covariance matrices of genetic and residual effects were set as

$$\mathrm{cov}(\mathbf{g_0}, \mathbf{g_1}, \boldsymbol{\gamma}) = \begin{bmatrix} \sigma^2_{\mathbf{g_0}} & \sigma_{\mathbf{g_0},\mathbf{g_1}} & \sigma_{\mathbf{g_0},\boldsymbol{\gamma}} \\ \sigma_{\mathbf{g_0},\mathbf{g_1}} & \sigma^2_{\mathbf{g_1}} & \sigma_{\mathbf{g_1},\boldsymbol{\gamma}} \\ \sigma_{\mathbf{g_0},\boldsymbol{\gamma}} & \sigma_{\mathbf{g_1},\boldsymbol{\gamma}} & \sigma^2_{\boldsymbol{\gamma}} \end{bmatrix} = \begin{bmatrix} (1-\sigma^2_{\mathbf{g_1}}-\sigma^2_{\boldsymbol{\tau}_1})/2 & 0 & \sigma_{\mathbf{g_0},\boldsymbol{\gamma}} \\ 0 & 0.1 & 0 \\ \sigma_{\mathbf{g_0},\boldsymbol{\gamma}} & 0 & 0.5 \end{bmatrix}$$

and

$$\mathrm{cov}(\boldsymbol{\tau_0}, \boldsymbol{\tau_1}, \boldsymbol{\varepsilon}) = \begin{bmatrix} \sigma^2_{\boldsymbol{\tau_0}} & \sigma_{\boldsymbol{\tau_0},\boldsymbol{\tau_1}} & \sigma_{\boldsymbol{\tau_0},\boldsymbol{\varepsilon}} \\ \sigma_{\boldsymbol{\tau_0},\boldsymbol{\tau_1}} & \sigma^2_{\boldsymbol{\tau_1}} & \sigma_{\boldsymbol{\tau_1},\boldsymbol{\varepsilon}} \\ \sigma_{\boldsymbol{\tau_0},\boldsymbol{\varepsilon}} & \sigma_{\boldsymbol{\tau_1},\boldsymbol{\varepsilon}} & \sigma^2_{\boldsymbol{\varepsilon}} \end{bmatrix} = \begin{bmatrix} (1-\sigma^2_{\mathbf{g_1}}-\sigma^2_{\boldsymbol{\tau}_1})/2 & 0 & \sigma_{\boldsymbol{\tau_0},\boldsymbol{\varepsilon}} \\ 0 & 0 & 0 \\ \sigma_{\boldsymbol{\tau_0},\boldsymbol{\varepsilon}} & 0 & 0.5 \end{bmatrix}$$

where $\sigma^2_{\mathbf{g_0}}$ and $\sigma^2_{\boldsymbol{\tau_0}}$ were $(1-\sigma^2_{\mathbf{g_1}}-\sigma^2_{\boldsymbol{\tau_1}})/2$, therefore, these were set as 0.45 in the presence of $\sigma^2_{\mathbf{g_0}}$=0.1 without RxE effects, and $\sigma_{\mathbf{g_0},\boldsymbol{\gamma}}$ and $\sigma_{\boldsymbol{\tau_0},\boldsymbol{\varepsilon}}$ are varied from 0 to 0.1 in accordance with the simulation scenario.

In the presence of both GxE and RxE effects (denoted as full scenario), the variance and covariance matrix for genetic and residual effects were set as

$$\mathrm{cov}(\mathbf{g_0}, \mathbf{g_1}, \boldsymbol{\gamma}) = \begin{bmatrix} \sigma^2_{\mathbf{g_0}} & \sigma_{\mathbf{g_0},\mathbf{g_1}} & \sigma_{\mathbf{g_0},\boldsymbol{\gamma}} \\ \sigma_{\mathbf{g_0},\mathbf{g_1}} & \sigma^2_{\mathbf{g_1}} & \sigma_{\mathbf{g_1},\boldsymbol{\gamma}} \\ \sigma_{\mathbf{g_0},\boldsymbol{\gamma}} & \sigma_{\mathbf{g_1},\boldsymbol{\gamma}} & \sigma^2_{\boldsymbol{\gamma}} \end{bmatrix} = \begin{bmatrix} (1-\sigma^2_{\mathbf{g_1}}-\sigma^2_{\boldsymbol{\tau}_1})/2 & 0 & \sigma_{\mathbf{g_0},\boldsymbol{\gamma}} \\ 0 & 0.1 & 0 \\ \sigma_{\mathbf{g_0},\boldsymbol{\gamma}} & 0 & 0.5 \end{bmatrix}$$

and

$$\mathrm{cov}(\boldsymbol{\tau_0}, \boldsymbol{\tau_1}, \boldsymbol{\varepsilon}) = \begin{bmatrix} \sigma^2_{\boldsymbol{\tau_0}} & \sigma_{\boldsymbol{\tau_0},\boldsymbol{\tau_1}} & \sigma_{\boldsymbol{\tau_0},\boldsymbol{\varepsilon}} \\ \sigma_{\boldsymbol{\tau_0},\boldsymbol{\tau_1}} & \sigma^2_{\boldsymbol{\tau_1}} & \sigma_{\boldsymbol{\tau_1},\boldsymbol{\varepsilon}} \\ \sigma_{\boldsymbol{\tau_0},\boldsymbol{\varepsilon}} & \sigma_{\boldsymbol{\tau_1},\boldsymbol{\varepsilon}} & \sigma^2_{\boldsymbol{\varepsilon}} \end{bmatrix} = \begin{bmatrix} (1-\sigma^2_{\mathbf{g_1}}-\sigma^2_{\boldsymbol{\tau}_1})/2 & 0 & \sigma_{\boldsymbol{\tau_0},\boldsymbol{\varepsilon}} \\ 0 & 0.1 & 0 \\ \sigma_{\boldsymbol{\tau_0},\boldsymbol{\varepsilon}} & 0 & 0.5 \end{bmatrix}$$

where $\sigma^2_{\mathbf{g_1}}$ and $\sigma^2_{\boldsymbol{\tau_1}}$ were 0.1, respectively, therefore, $\sigma^2_{\mathbf{g_0}}$ and $\sigma^2_{\boldsymbol{\tau_0}}$, which are $(1-\sigma^2_{\mathbf{g_1}}-\sigma^2_{\boldsymbol{\tau_1}})/2$, were set as 0.4, and $\sigma_{\mathbf{g_0},\boldsymbol{\gamma}}$ and $\sigma_{\boldsymbol{\tau_0},\boldsymbol{\varepsilon}}$ (genetic and residual correlations between the main trait and environment) were considered from 0 to 0.2 in accordance with the simulation scenario.

#### Binary disease traits

For binary disease simulations, the same ARIC data and simulation procedures were used as in the simulation of quantitative traits except that the simulated quantitative phenotypes were converted to the binary responses according to the population prevalence using a liability threshold model. We considered the population prevalence k=0.025, 0.05, 0.1 or 0.5. For example, we assigned affected or unaffected status to individuals with their standardised quantitative phenotypes (i.e. liability) greater or lower than the threshold of the standardised normal distribution truncating the proportion of k. It is noted that the threshold truncating the proportion k=0.025, 0.05, 0.1 or 0.5 is 1.96, 1.64, 1.28 or 0.

Similar to the simulations of quantitative traits, various scenarios mimicking real situations were tested in these binary disease simulations. The variance and covariance matrix of genetic effects on the liability scale was set as

$$\mathrm{cov}(\mathbf{g_0}, \mathbf{g_1}, \boldsymbol{\gamma}) = \begin{bmatrix} \sigma^2_{\mathbf{g_0}} & \sigma_{\mathbf{g_0},\mathbf{g_1}} & \sigma_{\mathbf{g_0},\boldsymbol{\gamma}} \\ \sigma_{\mathbf{g_0},\mathbf{g_1}} & \sigma^2_{\mathbf{g_1}} & \sigma_{\mathbf{g_1},\boldsymbol{\gamma}} \\ \sigma_{\mathbf{g_0},\boldsymbol{\gamma}} & \sigma_{\mathbf{g_1},\boldsymbol{\gamma}} & \sigma^2_{\boldsymbol{\gamma}} \end{bmatrix} = \begin{bmatrix} (0.5 - \sigma^2_{\mathbf{g_1}}) & 0 & 0 \\ 0 & \sigma^2_{\mathbf{g_1}} & 0 \\ 0 & 0 & 0.5 \end{bmatrix}$$

where $\sigma^2_{\mathbf{g_0}}$ and $\sigma^2_{\mathbf{g_1}}$ are the main genetic and GxE variances on the liability scale and they were simulated such that $\sigma^2_{\mathbf{g_1}}$ varied from 0 to 0.1 with $\sigma^2_{\mathbf{g_0}}$ = 0.5 - $\sigma^2_{\mathbf{g_1}}$. Therefore, the main genetic variance would be $\sigma^2_{\mathbf{g_0}}$= 0.5 in the absence of GxE (e.g. null or RxE only scenario). We further tested scenarios in the presence of the GxE with $\sigma^2_{\mathbf{g_1}}$= 0.01, 0.025, 0.05 and 0.1 (therefore, $\sigma^2_{\mathbf{g_0}}$=0.49, 0.475, 0.45 and 0.4). The variance and covariance matrix of residual effects was set as

$$\mathrm{cov}(\boldsymbol{\tau_0}, \boldsymbol{\tau_1}, \boldsymbol{\varepsilon}) = \begin{bmatrix} \sigma^2_{\boldsymbol{\tau_0}} & \sigma_{\boldsymbol{\tau_0},\boldsymbol{\tau_1}} & \sigma_{\boldsymbol{\tau_0},\boldsymbol{\varepsilon}} \\ \sigma_{\boldsymbol{\tau_0},\boldsymbol{\tau_1}} & \sigma^2_{\boldsymbol{\tau_1}} & \sigma_{\boldsymbol{\tau_1},\boldsymbol{\varepsilon}} \\ \sigma_{\boldsymbol{\tau_0},\boldsymbol{\varepsilon}} & \sigma_{\boldsymbol{\tau_1},\boldsymbol{\varepsilon}} & \sigma^2_{\boldsymbol{\varepsilon}} \end{bmatrix} = \begin{bmatrix} (0.5 - \sigma^2_{\boldsymbol{\tau_1}}) & 0 & 0 \\ 0 & \sigma^2_{\boldsymbol{\tau_1}} & 0 \\ 0 & 0 & 0.5 \end{bmatrix}$$

where $\sigma^2_{\boldsymbol{\tau_0}}$ and $\sigma^2_{\boldsymbol{\tau_1}}$ are the residual and RxE variances on the liability scale and they were simulated such that $\sigma^2_{\boldsymbol{\tau_1}}$ varied from 0 to 0.1 with $\sigma^2_{\boldsymbol{\tau_0}}$=0.5 - $\sigma^2_{\boldsymbol{\tau_1}}$. Therefore, the residual variance would be $\sigma^2_{\boldsymbol{\tau_0}}$=0.5 in the absence of RxE (e.g. null or GxE only scenario). We further tested scenarios in the presence of the RxE with $\sigma^2_{\boldsymbol{\tau_1}}$=0.01, 0.025, 0.05 and 0.1 (therefore, $\sigma^2_{\boldsymbol{\tau_0}}$=0.49, 0.475, 0.45 and 0.4).

#### Simulation with collider bias

The phenotypic data with collider bias were simulated to check whether the type I error rate of GxEsum was inflated or not. For this simulation, the null or RxE only scenario, i.e. in the absence of GxE effects, was used, in which the GxE variance was simulated as $\sigma^2_{\mathbf{g_1}}$ = 0, and the RxE variance was simulated as $\sigma^2_{\tau_1}$ = 0 or 0.1. In the null scenario, the main genetic and residual variances were $\sigma^2_{\mathbf{g_0}}$ = $\sigma^2_{\tau_0}$ = 0.5, and in the RxE only scenario, they were $\sigma^2_{\mathbf{g_0}}$ = $\sigma^2_{\tau_0}$ = 0.45.

Following Munafò et al. (2018) [1] and Yu et al. (2020) [2], the participation probability for each individual with the main trait and environment can be expressed as

$$\mathbf{p} = \frac{1}{1 + \exp\left(-\ln\left(OR_{\mathbf{y}}\right) * \mathbf{y} - \ln(OR_{\mathbf{c}}) * \mathbf{c}\right)}$$

where **p** is the vector of participation probabilities for individuals in a study, and **y** and **c** indicate the phenotypes of the main trait and the environmental variable. The selection odds ratios for the main trait and environment were notated as $OR_{\mathbf{y}}$ and $OR_{\mathbf{c}}$, respectively, which were set as 1.5 and 2 in our simulations. Therefore, individual's participation probability is determined by the joint effects of **y** and **c,** which generates a spurious association between y and c.

For quantitative trait, individuals with **p** smaller than a random number from the standardized uniform distribution (i.e. ~ U(0, 1)) were removed. After selection based on the collider model, about 3,600 individuals which were half of the total samples remained. For binary traits, the quantitative phenotypes were converted to the binary format based on a population prevalence k=0.1. Then, the same procedures to estimate **p** with which individuals were selected out.

#### 3. Theory Verification

To verify our proposed model, the observed intercept values from all simulations were compared to the theoretical values, which are expected as close to 1 for the main genetic variance (g0) [3] and 1+2$h^2_{g_1}$+ 2$h^2_{\tau_1}$ for the interaction variance (g1) when $\mathbf{X}_j$ is normally distributed. $h^2_{g_1}$ and $h^2_{\tau_1}$ indicate the proportion of phenotypic variance explained by GxE and RxE. When the environmental variable (**E**) is non-normal, the expected intercept can be expressed as $1 + (k_{urtosis}-1)h^2_{\boldsymbol{g_1}} + (k_{urtosis}-1)h^2_{\boldsymbol{\tau_1}}$, where $k_{urtosis}$ indicates the kurtosis of **E**. The expected intercept values from the theory (i.e. eq. (4) and eq. (5)) are agreed with the observed values. In further simulations with additional confounding effects such as G-E and/or R-E correlations, the expected and observed values have an excellent agreement (Supplementary Table 2).

#### 4. Details of data obtained from UK Biobank

##### Main traits

For the quantitative trait, we used BMI (UK Biobank data field: 21001) that is calculated as weight/height. There were 287,923 non-missing records out of 288,837 individuals. For the binary disease trait, we used hypertension. Following the previous study using the UK Biobank participants with hypertension, we collected the patient information who were having systolic blood pressure (UK Biobank data field: 4080) ≥ 140 mmHg, or diastolic blood pressure (UK Biobank data field: 4079) ≥ 90 mmHg, or who had prescribed medication for blood pressure (UK Biobank data field: 6177). Each blood pressure was measured twice at a slight time difference, and we used the mean of these two measurements to obtain the binary disease trait. As a result, we obtained the UK biobank participants with hypertension with a population prevalence of 0.51. There were 266,144 non-missing records out of 288,837 individuals.

##### Covariates

For the quantitative trait analyses, we used 4 environmental variables in the GxE model to estimate how much genetic effects of BMI were modulated by these environmental variables. The environmental variables consist of neuroticism score (NEU), alcohol intake frequency, physical activity and age. NEU was obtained from the summary score of neuroticism that was collected from answers by patients about 12 domains of neurotic behaviours. Alcohol intake frequency (UK Biobank data field: 1558) was obtained from asking “How often do you drink alcohol”. Individuals who preferred not to answer were coded as missing. We used the summed metabolic equivalent task (MET) minutes per week as the physical activity variable. We also used age of participant at the recruitment as the environmental variable in the model.

For the binary disease analysis, we used 3 variables for the environments in the GxE model to estimate the extent to which genetic effects of hypertension were modulated by these environments. These variables are BMI, WHR and BFP. BMI was used as the main trait in the quantitative trait analysis but used as environment in the binary trait analysis. WHR is calculated as waist circumference (UK Biobank data field: 48) divided by hip measurements (UK Biobank data field: 49), and the BFP (UK Biobank data field: 23099) was obtained from the impedance results.

We also used type 2 diabetes as another example application of GxEsum with 3 environmental variables, i.e. BMI, and diastolic and systolic blood pressures (BP) that are known to be risk factors. BP measurement was performed twice after the participant had been at rest for 5 minutes, and the mean of two measures for systolic (UK Biobank data field: 4079) or diastolic (UK Biobank data field 4080) was used in this study.

##### Confounders

The main traits, which are BMI and patients with hypertension in this analysis, were pre-adjusted for confounders. Demographics, first 10 principal components and the environment which is fitting in the GxEsum model were considered as confounders, and the demographics consist of age at recruitment (UK Biobank data field: 21022), gender (UK Biobank data field: 31), assessment centre (UK Biobank data field: 54), Townsend deprivation index at recruitment (UK Biobank data field: 189), genotyping processed batch (UK Biobank data field: 22000), year of birth (UK Biobank data field: 34), average total household income before tax (UK Biobank data field: 738) and educational qualification (UK Biobank data field: 6138). Following the previous paper [4], the educational qualification was collected by considering all 6 responses (6138-0.0 – 6138-0.5) due to the possibility of multiple selections. Before considering the responses, we converted the score coded by UK Biobank to the continuous educational yearly measure and used the maximum value out of 6 variables for each individual.

#### 5. Demonstration of GxEsum software

GxEsum shell script and example files can be downloaded from https://github.com/honglee0707/GxEsum

README file instruct how to run GxEsum as in the following.

#### Users should prepare LDSC (munge_sumstats.py and ldsc.py) and PLINK (v1.9) software that are ready to run in the folder.

* if LDSC is not working, please try to upgrade python pacakge, e.g. pip install --upgrade numpy==1.16.0
(change numpy==1.12 to numpy==1.16.0 in the environment.yml file for using Anaconda3. You can fix the environment.yml file)

### GWAS process

: using plink, GWAS summary stats including the GxE component can be obtained (BETA by fitting to the linear regression model).

e.g., ./plink1.9 --bfile example --linear interaction --pheno example_quant.pheno --covar example.cov --parameters 1, 2, 3 --allow-no-sex --out example

or for the phenotype formatted as binary,

e.g., ./plink1.9 --bfile example --linear interaction --pheno example_binary.pheno --covar example.cov --parameters 1, 2, 3 --allow-no-sex --out example
(for binary traits, we have assigned as 20/10 to fit into the linear regression model)

-> This plink command will give "example.assoc.linear" ( contains $BETA for regression coefficients)

Note. Using different versions of PLINK (e.g., plink2) will give the different format of outcomes, users should modify the R script depending on which version of plink is used.

### Converting your GWAS summary statistics to LDSC format (e.g., .sumstats.gz)

For LDSC, the outcomes of GWAS needs to be converted to .sumstats.format (see LDSC instruction)
They strongly suggest that using ./munge_sumstats.py which is included with ldsc.

Before reformatting the data, an extra process for munging your summary statisitcs is required.

I would recommend using the R script (see input_for_munge.R) that can help preparing the input file for munge.

The input_for_munge.R is a script for extracting the information of GxE effects which is required in munging process.

The command in shell script to process the R script is

e.g., R CMD BATCH --no-save input_for_munge.R

### Munge your data using the generated input file from R script above

e.g., ./munge_sumstats.py --sumstats example.ldsc --merge-alleles w_hm3.snplist --out example

This command will output .sumstats.gz format file that is generated based on your GWAS summary statistics

* To make munge_sumstats.py complete GWAS Summary statistics conversion faster, you can reduce the chunksize from 5000000 (default) to 500000 by adding the option --chunksize 500000.

### Estimating the phenotypic variance explained by GxE effects (LD score regression)

using the LD Score that can be estimated based on your own plink binary data or downloaded the estimated 1000 Genomes Europeans LDSC.

e.g., ./ldsc.py --h2 example.sumstats.gz --ref-ld ldsc_example/example --w-ld ldsc_example/example --out example

This will give .log file that contains the phenotypic variance explained by GxE and intercept value.

Note. It is recommended using LD Scores estimated based on in-sample if individual-level genotypes are available.

When running GxEsum_master.sh, the output should be

```
*********************************************************
* GxEsum
* Jisu Shin & S. Hong Lee (2020)
```

```
* University of South Australia
*********************************************************

* GWAS for linear regression model
* done

* LDSC for GxE interaction, i.e. var(g1)
* done

* LDSC for main genetic effects, i.e. var(g0)
done

* GxEsum analysis results
                       Main genetic effects      GxE interaction
Estimated intercept  :    0.9109                 1.1186
Estimated variance   :    4.1696                 1.1557
SE                   :   3.5271                  3.91
P-value              :   0.2371418               0.7675543

* NOTE
  The total phenotypic variance is assumed to be 1
  P-value is from Wald test
  This toy example is just to demonstrate how to run GxEsum, which can generate
unreliable estimates with large SE (a user may need larger # samples and SNPs)
```

#### Supplementary Figures

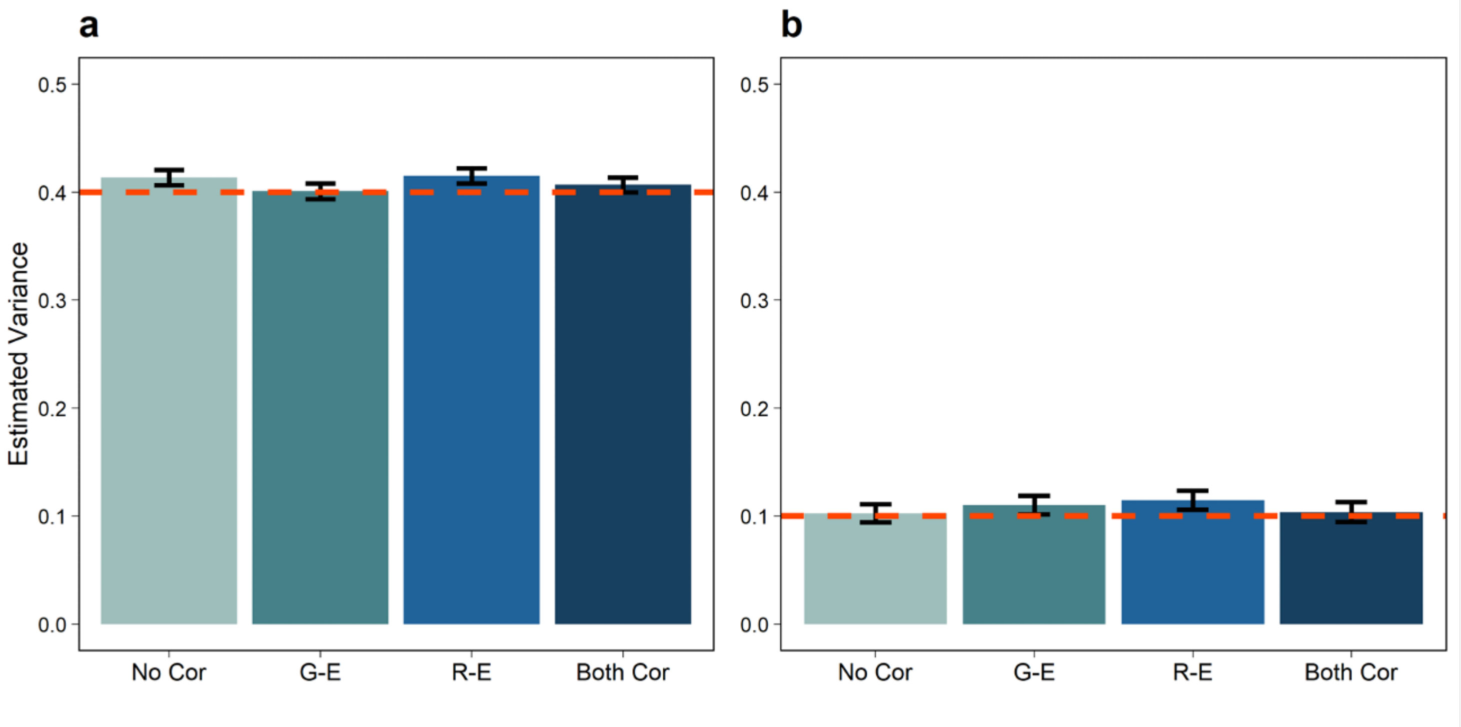


**Supplementary Figure 1. Estimated variance components of main genetic (a) and GxE effects (b) using GxEsum in the presence of GxE and RxE (i.e. full scenario)** The true simulated values are marked as dashed horizontal lines (0.4 and 0.1 for the main genetic and GxE variances). The bars and vertical error bars indicate the estimated variances and 95% confidence intervals (CI) that were obtained from 500 replicates. The environmental variable was simulated as $E = \gamma + \varepsilon$ where $\gamma$ is the genetic effects and $\varepsilon$ is residual with var($\gamma$)=0.5 and var($\varepsilon$)=0.5 such that heritability of the environmental variable is 0.5 (see Supplementary Note 2).

No Cor: There is no G-E or R-E correlation.
G-E: There is G-E correlation only (the genetic correlation between the trait and environment is set as 0.1).
R-E: There is R-E correlation only (the residual correlation between the trait and environment is set as 0.1).
Both Cor: Both G-E and R-E correlations exist (the genetic and residual correlation between the trait and environment is set as 0.1, respectively).

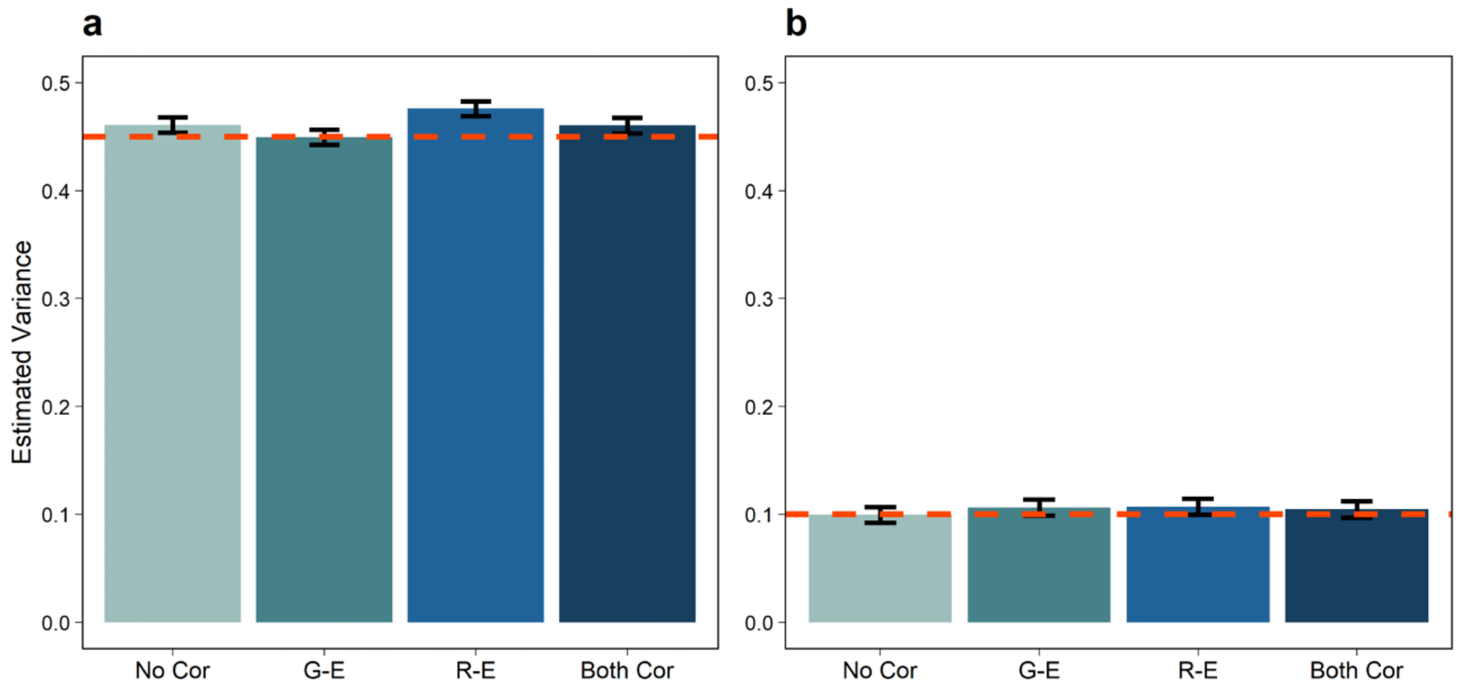


**Supplementary Figure 2. Estimated variance components of main genetic (a) and GxE effects (b) using GxEsum in the presence of GxE without RxE (i.e. GxE only scenario).** The true simulated values are marked as dashed horizontal lines (0.45 and 0.1 for the main genetic and GxE variances). The bars and vertical error bars indicate the estimated variances and 95% CI that were obtained from 500 replicates. The environmental variable was simulated as $E = \gamma + \varepsilon$ where $\gamma$ is the genetic effects and $\varepsilon$ is residual with var($\gamma$)=0.5 and var($\varepsilon$)=0.5 such that heritability of the environmental variable is 0.5.

No Cor: There is no G-E and R-E correlation
G-E: There is G-E correlation only (the genetic correlation between the main trait and environment is set as 0.1).
R-E: There is R-E correlation only (the residual correlation between the main trait and environment is set as 0.1).
Both Cor: Both G-E and R-E correlations exist (the genetic and residual correlation between the main trait and environment is set as 0.1, respectively).

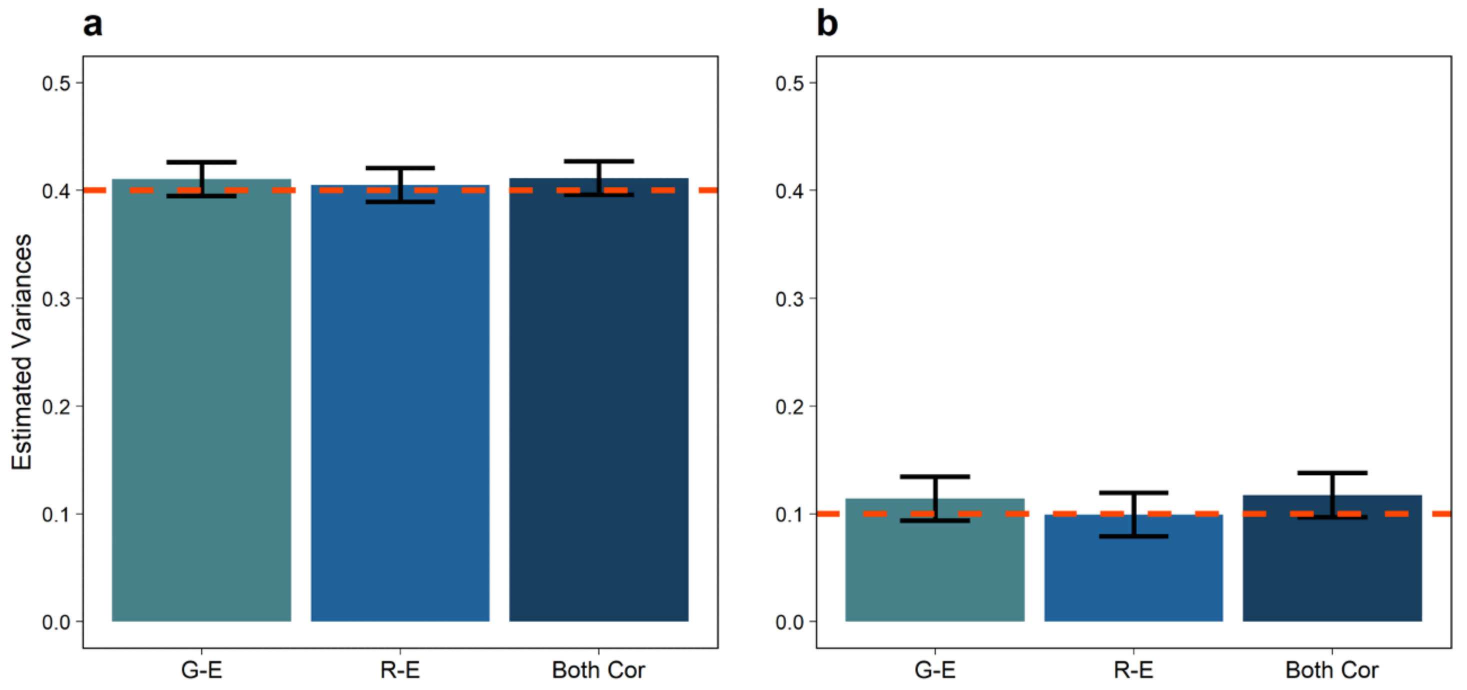


**Supplementary Figure 3. Estimated variance components of main genetic (a) and GxE effects (b) using GxEsum when the possible correlations are smaller effects in the presence of both interactions (i.e. full scenario).** The true simulated values are marked as dashed horizontal lines (0.4 and 0.1 for the main genetic and GxE variances). The error bars indicate 95% CI from 100 replicates. The environmental variable was simulated as $E = \gamma + \varepsilon$ where $\gamma$ is the genetic effects and $\varepsilon$ is residual with var($\gamma$)=0.5 and var($\varepsilon$)=0.5 such that heritability of the environmental variable is 0.5.

G-E: There is G-E correlation only (the genetic correlation between the main trait and environment is set as 0.05).
R-E: There is R-E correlation only (the genetic correlation between the main trait and environment is set as 0.05).
Both Cor: Both G-E and R-E correlations exist (the genetic and residual correlation between the main trait and environment is set as 0.05, respectively).

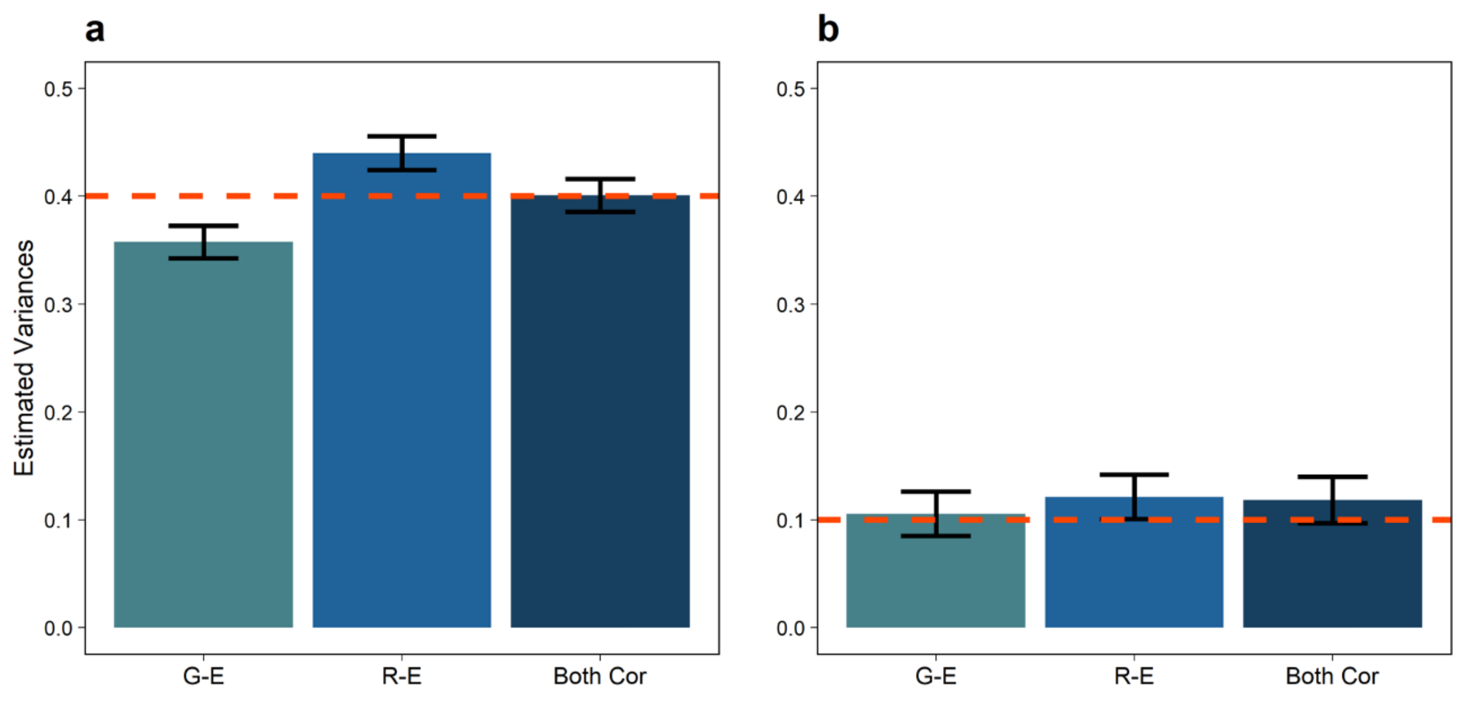


**Supplementary Figure 4. Estimated variance components of main genetic (a) and GxE effects (b) using GxEsum when the possible correlations are larger effects in the presence of both interactions (i.e. full scenario).** The true simulated values are marked as dashed horizontal lines (0.4 and 0.1 for the main genetic and GxE variances). The error bars indicate 95% CI from 100 replicates. The environmental variable was simulated as $E = \gamma + \varepsilon$ where $\gamma$ is the genetic effects and $\varepsilon$ is residual with var($\gamma$)=0.5 and var($\varepsilon$)=0.5 such that heritability of the environmental variable is 0.5.

G-E: There is G-E correlation only (the genetic correlation between the trait and environment is set as 0.2).
R-E: There is R-E correlation only (the residual correlation between the trait and environment is set as 0.2).
Both Cor: Both G-E and R-E correlations exist (the genetic and residual correlation between the trait and environment is set as 0.2, respectively).

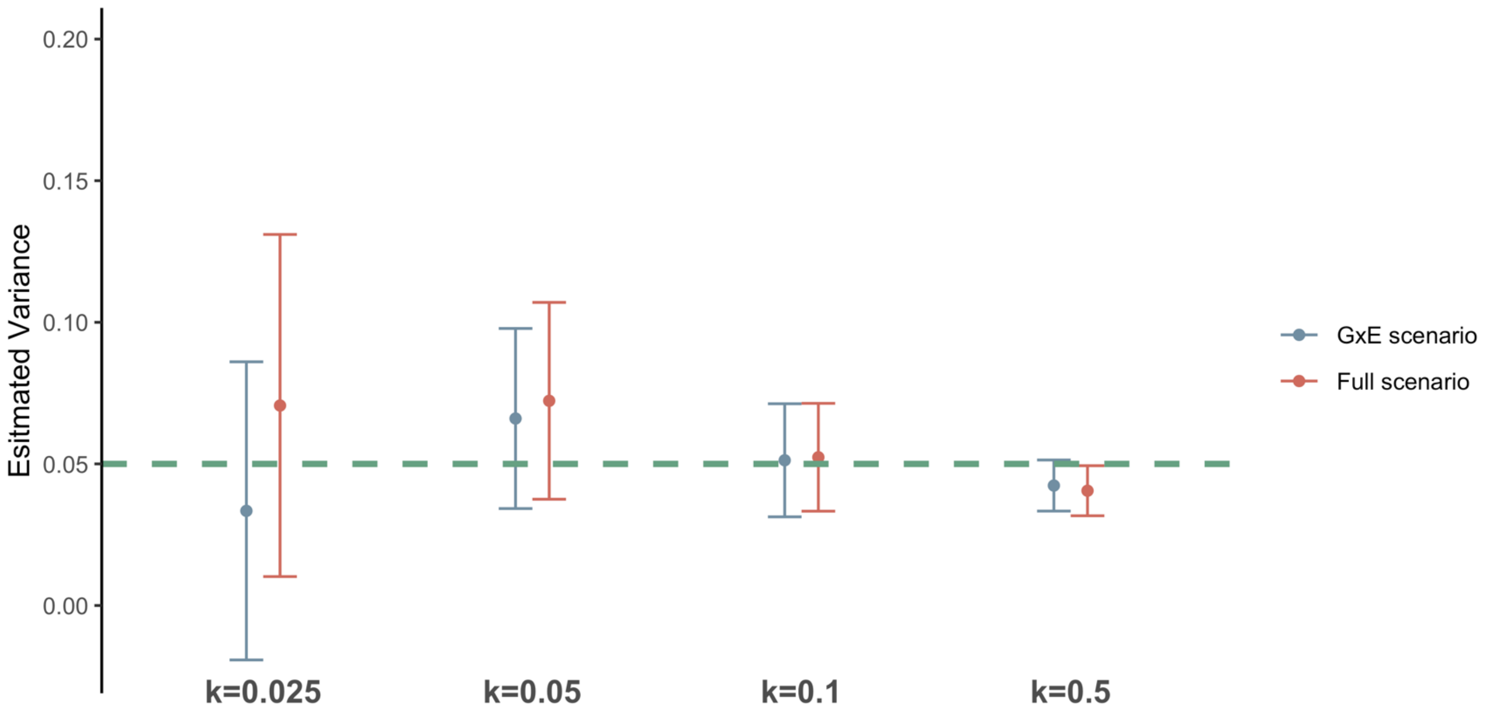


**Supplementary Figure 5. Estimated variance component of GxE when binary disease traits were used.** The GxE was estimated under the scenarios with different prevalence rates, which are k=2.5%, 5%, 10%, 50%. The point indicates the mean of variance and vertical error bar is 95% CI that were obtained from the average of 500 replicates, and the green horizontal dashed line is the true GxE value that is set as 0.05 in the scenarios. Estimates marked as blue and red colour is obtained from GxE only and full model, respectively.

GxE scenario: Simulation with GxE only ($\sigma^2_{\mathbf{g_1}}$ =0.05).
Full scenario: Simulation with GxE ($\sigma^2_{\mathbf{g_1}}$= 0.05) and RxE ($\sigma^2_{\boldsymbol{\tau_1}}$=0.05).

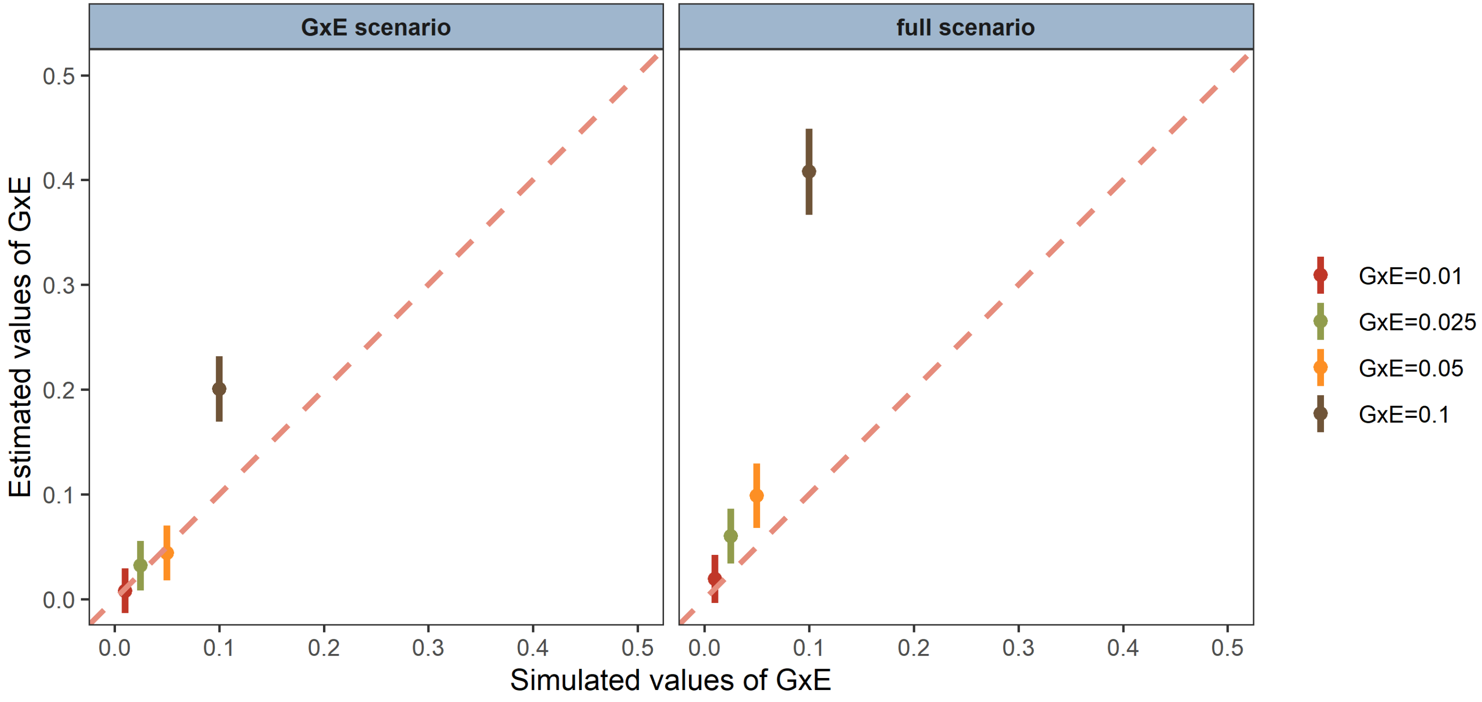


**Supplementary Figure 6. The level of biasness increases when the proportion of phenotypic variance explained by GxE increases in the case of population prevalence k=0.025.** The phenotypic variances explained by GxE were estimated under the GxE (left) and full (right) scenario with the different GxE variances, which are $\sigma^2_{\mathbf{g_1}}$=0.01, 0.025, 0.05 and 0.1 (noting that the phenotypic variance is 1). In the full scenario, the phenotypic variances explained by RxE ($\sigma^2_{\boldsymbol{\tau}_1}$) are the same values of the reflected GxE values. The error bars are 95% CI from 2000 replicates (which was required to correctly assess the biasness especially for the simulation of small GxE effects). The dashed diagonal line is where simulated and estimated values are equal.

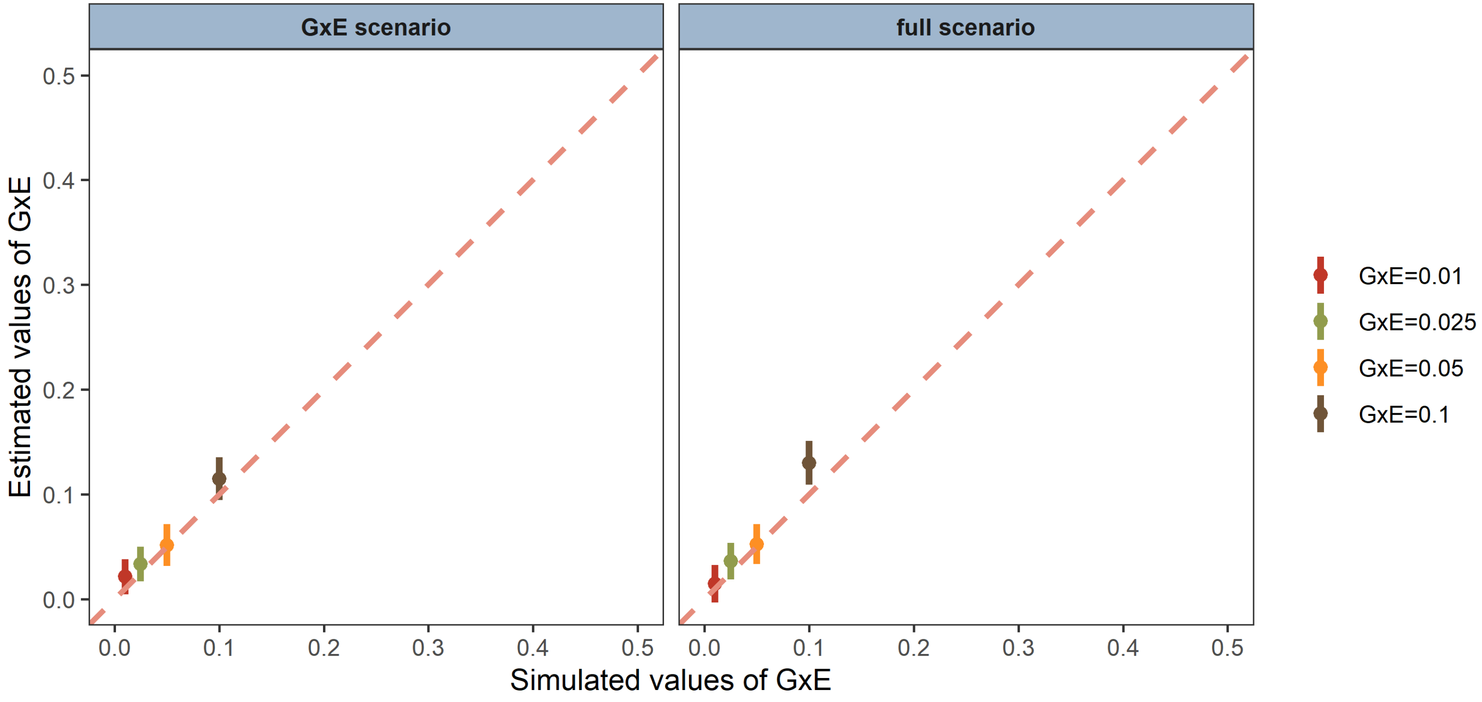


**Supplementary Figure 7. The level of biasness increases when the proportion of phenotypic variance explained by GxE increases in the case of population prevalence k=0.1.** The phenotypic variances explained by GxE were estimated under the GxE (left) and full (right) scenario with the different GxE variances, which are $\sigma^2_{\mathbf{g_1}}$=0.01, 0.025, 0.05 and 0.1 (noting that the phenotypic variance is 1). In the full scenario, the phenotypic variances explained by RxE ($\sigma^2_{\boldsymbol{\tau}_1}$) are the same values of the reflected GxE values. The error bars are 95% CI from 500 replicates and the dashed line is where simulated and estimated values are equal.

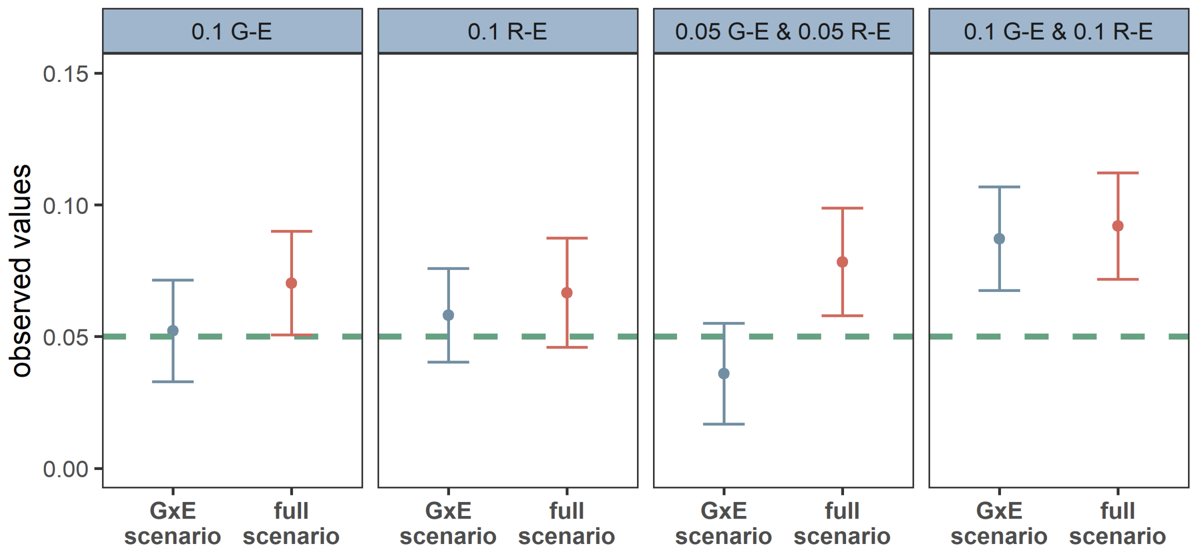


**Supplementary Figure 8. Estimated variance components of GxE when using binary disease traits (k=0.1) with which environmental variable is significantly correlated (i.e. G-E and/or R-E correlations).**

GxE (blue): Simulation with GxE only ($\sigma^2_{\mathbf{g}_1}$ =0.05). Full (red): Simulation with GxE ($\sigma^2_{\mathbf{g}_1}$ =0.05) and RxE ($\sigma^2_{\boldsymbol{\tau}_1}$=0.05). For GxE simulation or Full simulation model, additional simulations with G-E and/or R-E correlations were conducted to check how the confounders affected on the GxE estimation. There were four scenarios for the confounders with 1) a G-E correlation of 0.1 only (the first panel), 2) a R-E correlation of 0.1 only (the second panel), 3) both G-E and R-E correlations of 0.05 (the third panel) and 4) both G-E and R-E correlations of 0.1 (the last panel). The true simulated values (Var(GxE) =$\sigma^2_{\mathbf{g}_1}$=0.05) are marked as the green dashed horizontal line. The points and error bars indicate the estimates and 95% CI obtained from 500 replicates.

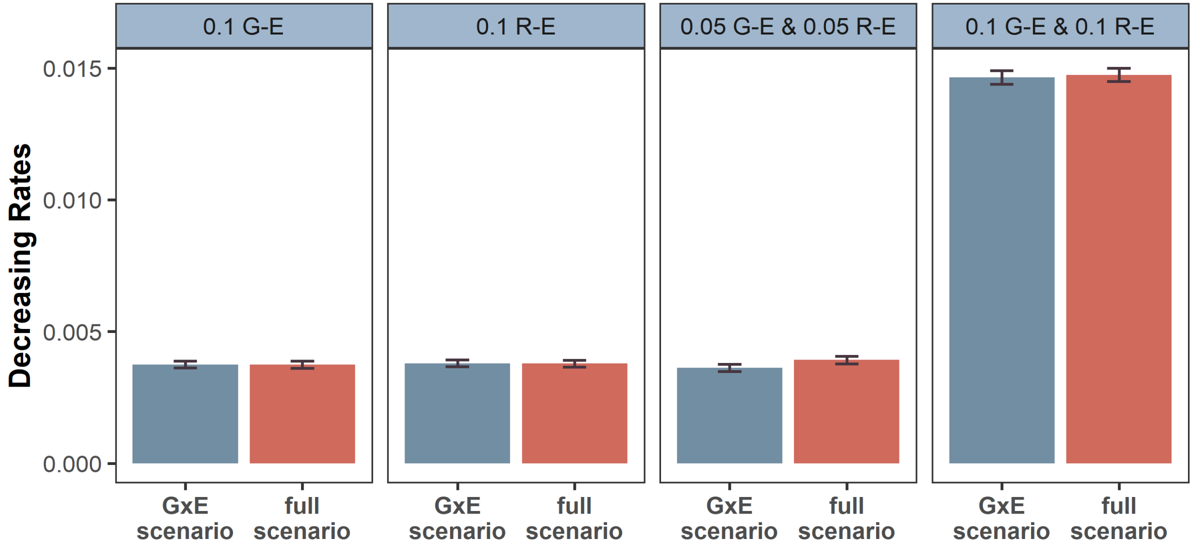


**Supplementary Figure 9. The decreasing rate of phenotypic variances after adjusting the phenotypes for the environment.** The ratio of phenotypic variance in the raw phenotypes to the pre-adjusted phenotypes was obtained in the presence of G-E or/and R-E correlations. The main bar indicates the mean of decreasing rates and error-bar represents the 95 % confidence interval of the mean decreasing rate averaged over 500 replicates. In each replicate, we simulated either GxE only (i.e. GxE scenario) or both GxE and RxE (i.e. full scenario), and 4 different G-E/R-E correlations were simulated and tested in each of the GxE scenario and full scenario, i.e. a G-E correlation of 0.1 (1$^{st}$ panel), a R-E correlation of 0.1 (2$^{nd}$ panel), both G-E and R-E correlations of 0.05 (3$^{rd}$ panel) and both G-E and R-E correlations of 0.1 (4$^{th}$ panel).

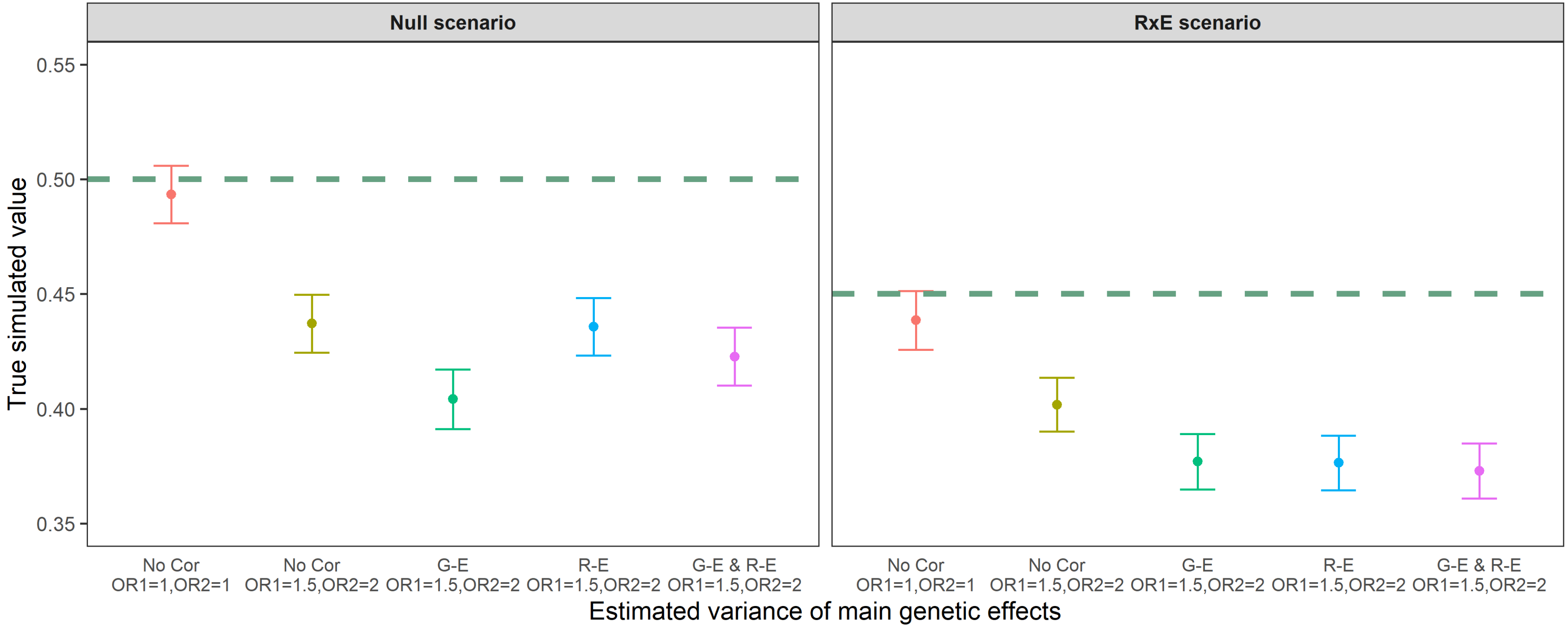


**Supplementary Figure 10. Estimated variance of main genetic effects using quantitative trait when the collider bias was considered.** The phenotypic variance explained by main genetic effects (i.e. SNP heritability, $\sigma^2_{g_0}$) was estimated under various scenarios, which are Null ($\sigma^2_{g_1}$=0 and $\sigma^2_{\tau_1}$=0) and RxE only ($\sigma^2_{g_1}$=0 and $\sigma^2_{\tau_1}$=0.1) scenarios. The green dashed line in each panel indicates the true simulated variance of main genetic effects that is 0.5 and 0.45, respectively, under the null and RxE scenario. Each point and error bar are the mean of the variance and 95% CI obtained from 500 replicates. These simulation results show that SNP heritability can be substantially underestimated because of collider bias (OR=1.5 and 2), which is not often observed in real data analyses even with a self-report study (e.g. UK Biobank). We arbitrarily set the bias factor with OR=1.5 and 2, which may be less in real situation (OR < 1.5). After selections with collider bias, the number of samples was around 3,600 individuals were used (see Supplementary Note 2).

No Cor: There is no correlation.
G-E: In the presence of G-E correlation
R-E: In the presence of R-E correlation
G-E & R-E: In the presence of both G-E and R-E correlations
OR1=1, OR2=1: The selection odd ratio was set as 1 and 1 for the main trait and environment, which is the same as the condition without the selection bias
OR1=1.5, OR2=2: The selection odd ratio was set as 1.5 and 2 for the main trait and environment.

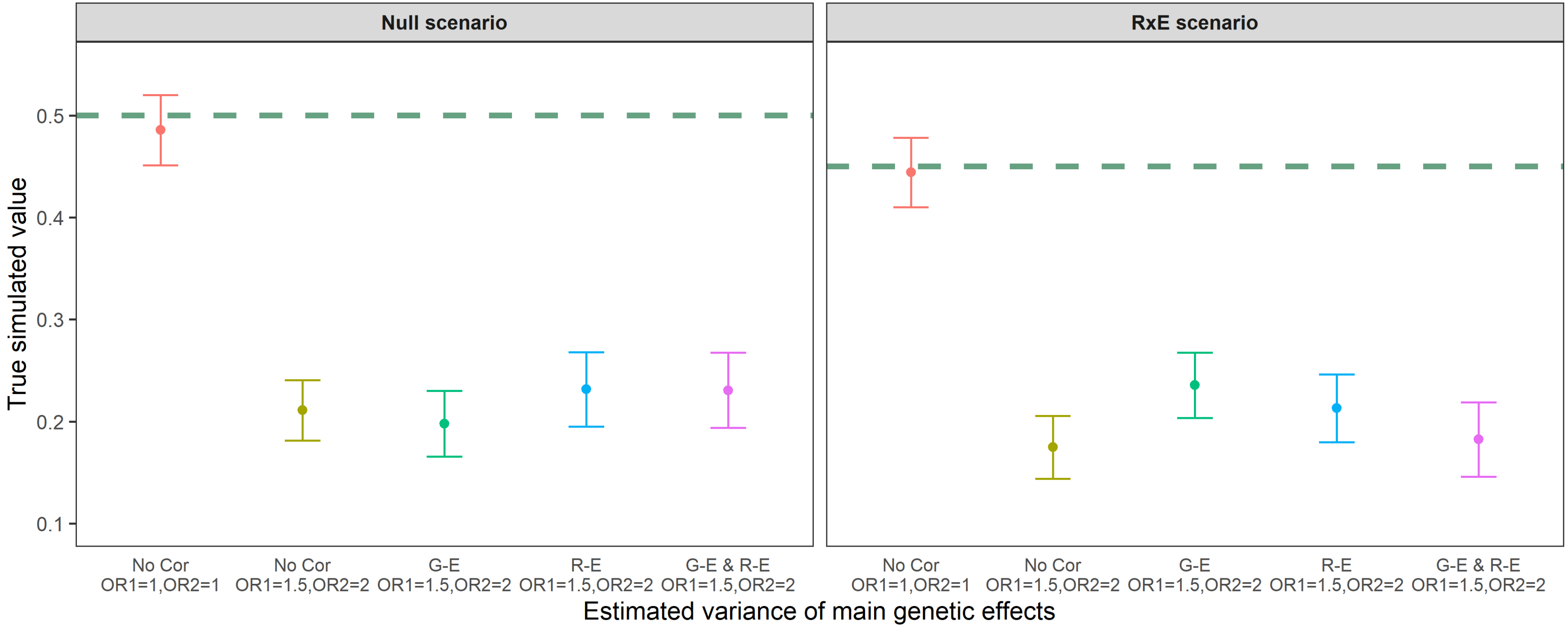


**Supplementary Figure 11. Estimated variance of main genetic effects using binary trait (k=0.1) when the collider bias was considered.** The phenotypic variance explained by main genetic effects (i.e. SNP heritability, $\sigma^2_{g_0}$) was estimated under various scenarios, which are Null ($\sigma^2_{g_1}$=0 and $\sigma^2_{\tau_1}$=0) and RxE only ($\sigma^2_{g_1}$=0 and $\sigma^2_{\tau_1}$=0.1) scenarios. The green dashed line in each panel indicates the true simulated variance of main genetic effects that is 0.5 and 0.45, respectively, under the null and RxE scenario. Each point and error bar are the mean of the variance and 95% CI obtained from 500 replicates. These simulation results show that SNP heritability can be substantially underestimated because of collider bias (OR=1.5 and 2), which is not often observed in real data analyses even with a self-report study (e.g. UK Biobank). We arbitrarily set the bias factor with OR=1.5 and 2, which may be less in real situation (OR <1.5). After selections with collider bias, the number of samples was around 3,600 individuals were used (see Supplementary Note 2). In comparison to the simulations using quantitative trait, it is noted that the usage of binary trait may cause more biased estimates rather than that of the quantitative traits when the range of collider bias equal.

No Cor: There is no correlation.
G-E: In the presence of G-E correlation
R-E: In the presence of R-E correlation
G-E & R-E: In the presence of both G-E and R-E correlations
OR1=1, OR2=1: The selection odd ratio was set as 1 and 1 for the main trait and environment, which is the same as the condition without the selection bias
OR1=1.5, OR2=2: The selection odd ratio was set as 1.5 and 2 for the main trait and environment.

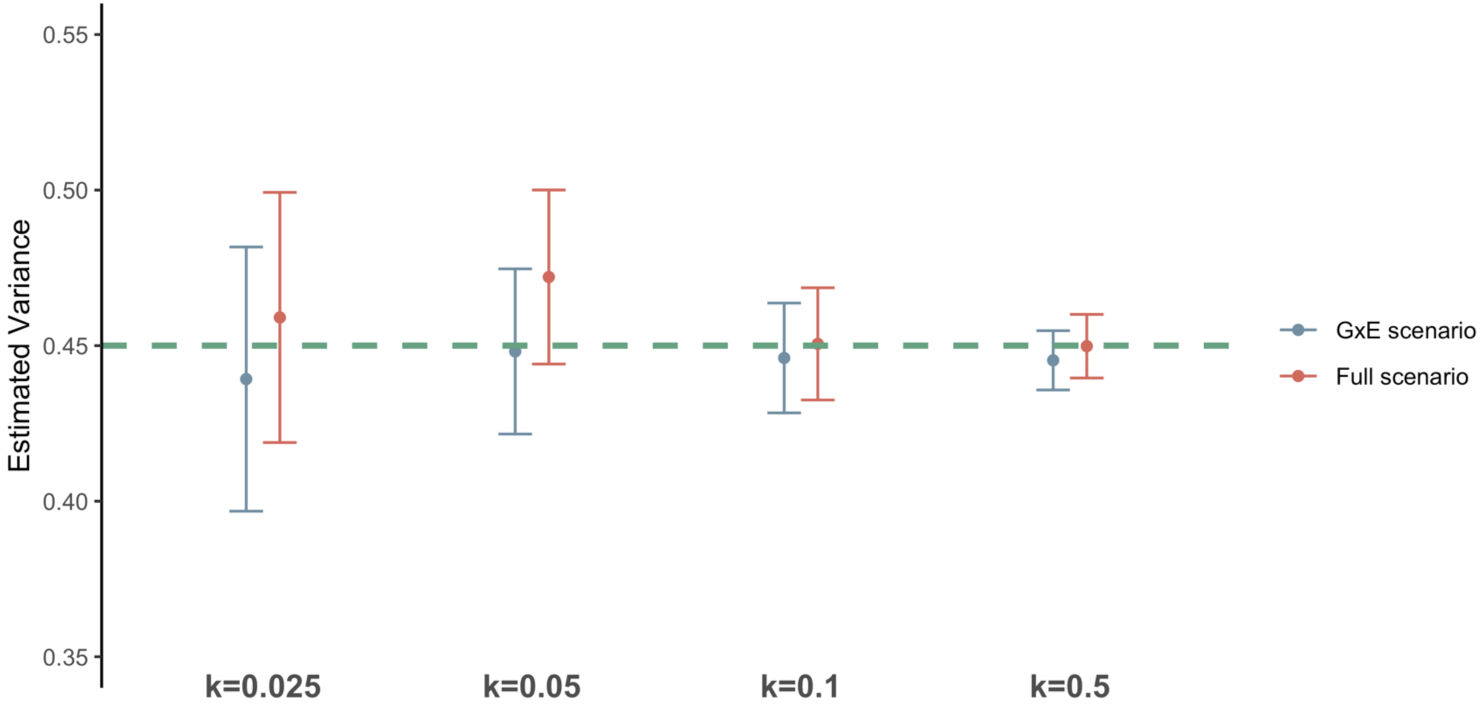


**Supplementary Figure 12. Estimated variance component of the main genetic effects (g0) when binary disease traits were used.** The phenotypic variances explained by the main genetic effects were estimated with different prevalence rates, which are k=2.5%, 5%, 10%, 50%. Each point and vertical error bar indicate the variance and 95% CI that were obtained from the average of 500 replicates, and the green horizontal dashed line is the true variance of main genetic effects that is set as 0.45 in scenarios. The results shown in red colour are obtained from simulations under the GxE only scenario, where the GxE is set as 0.05 without RxE, and the blue is obtained from the full scenario that is reflected both GxE ($\sigma^2_{\mathbf{g}_1}$=0.05) and RxE ($\sigma^2_{\boldsymbol{\tau}_1}$=0.05).

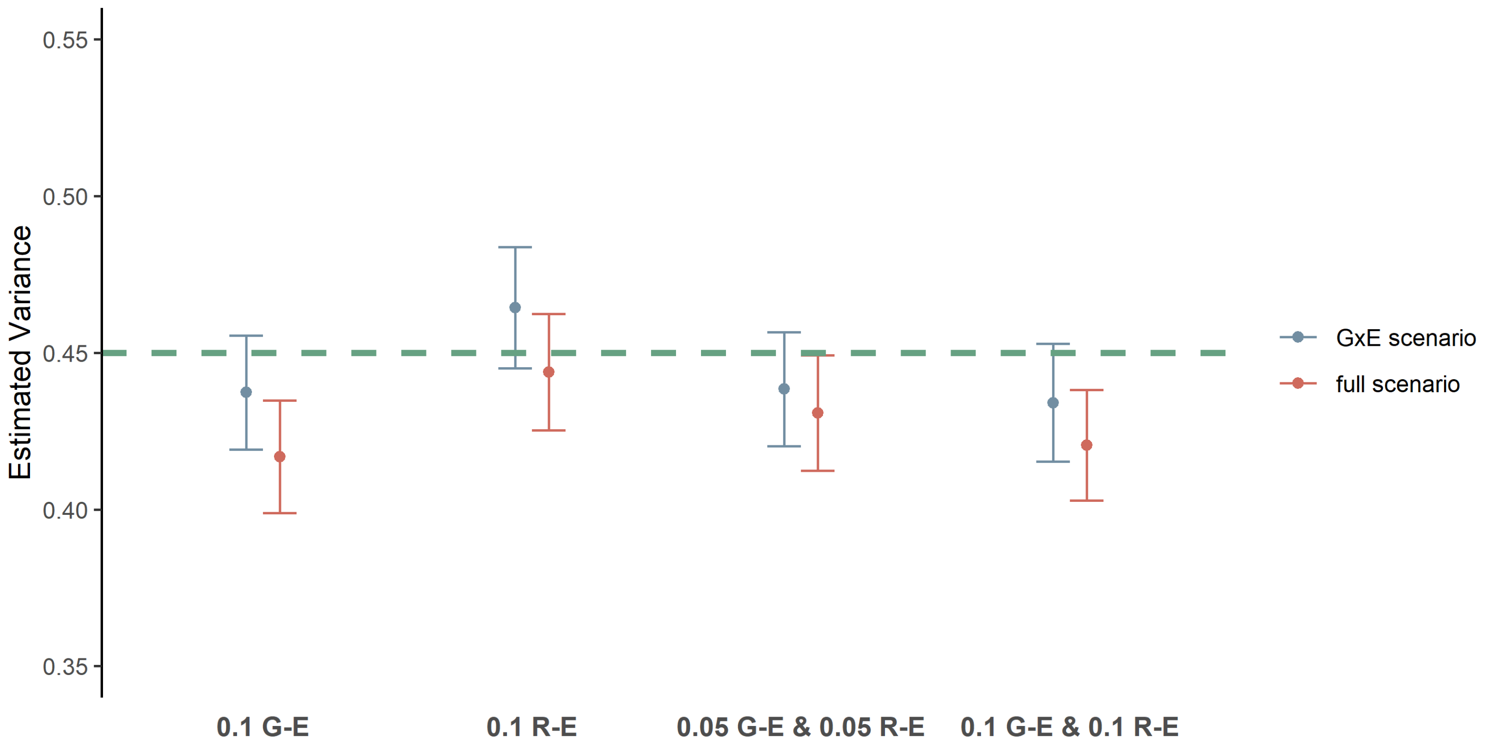


**Supplementary Figure 13. Estimated variance component of the main genetic effects (g0) when further potential confounders were considered in the case of population prevalence k=0.1.** Each point and vertical line indicate the estimate of heritability and 95 % CI that were observed from the simulations with potential confounders (e.g., G-E and/or R-E correlations), and the green dashed line is the true simulated heritability. In the GxE scenario (blue), the variances of GxE ($\sigma^2_{\mathbf{g_1}}$) was 0.05 without RxE ($\sigma^2_{\boldsymbol{\tau_1}}$), and the variances of GxE and RxE were respectively reflected as 0.05 for the full scenario (red).

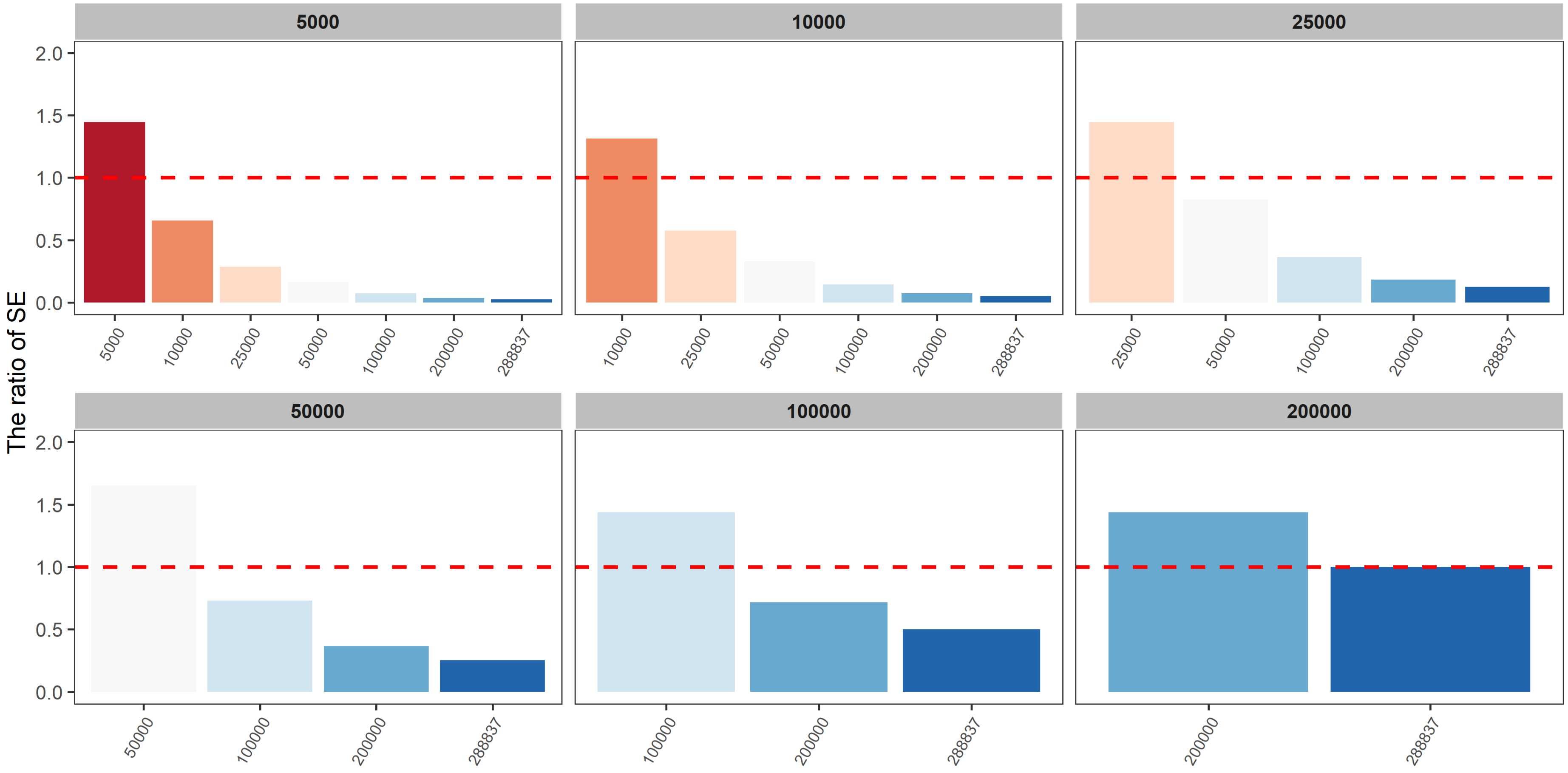


**Supplementary Figure 14. The ratio of SE from GxEsum to that from RNM using UK Biobank.** The ratio of SE in the usage of different sample sizes from 5,000 to 288,837 individuals were estimated, and the estimates were drawn into different panels under the usage of different sample sizes in theoretics. The x-axis of each panel shows the usage of different sample sizes in GxEsum analysis, and the y-axis that is the ratio of SE was obtained by dividing the values of GxEsum analysis into that of the theoretics. The bars indicate the ratio of SE, and the dashed red lines represent the ratio as 1.

### Supplementary Tables

##### Supplementary Table 1. Type I error rates of GxEsum when confounding effects are large.

| Scenarios | Type I Error rate |
|---|---|
| Var(GxE) = 0, Var(RxE) = 0 | 0.074 |
| Var(GxE) = 0, Var(RxE) = 0, G-E correlation = 0.25 | 0.03 |
| Var(GxE) = 0, Var(RxE) = 0, R-E correlation = 0.25 | 0.07 |
| Var(GxE) = 0, Var(RxE) = 0, G-E correlation=0.25, R-E correlation=0.25 | 0.068 |
| Var(GxE) = 0, Var(RxE) = 0.34 | 0.042 |
| Var(GxE) = 0, Var(RxE) = 0.34, G-E correlation = 0.2 | 0.036 |
| Var(GxE) = 0, Var(RxE) = 0.34, R-E correlation=0.2 | 0.042 |
| Var(GxE) = 0, Var(RxE) = 0.34, G-E correlation=0.2, R-E correlation=0.2 | 0.028 |
| **Average** | **0.048** |

**Supplementary Table 2. Theory verification using the obtained intercept values.**

| | | True value | | GxEsum analysis | |
|---|---|---|---|---|---|
| **Kurtosis=3 and skewness=0 (normal distribution) for environmental variable** | | | | | |
| **Model** | | $h^2_{g_1}$ | $h^2_{\tau_1}$ | **Expectation**[a] | **Observation**[b] |
| **No GxE & RxE** | No correlation | - | - | 1.0 | 0.998 (0.001) |
| | G-E correlation | - | - | 1.0 | 1.001 (0.001) |
| | E-E correlation | - | - | 1.0 | 0.999 (0.001) |
| | G-E & E-E correlations | - | - | 1.0 | 1.000 (0.001) |
| **GxE without RxE** | No correlation | 0.1 | - | 1.2 | 1.204 (0.002) |
| | G-E correlation | 0.1 | - | 1.2 | 1.202 (0.001) |
| | E-E correlation | 0.1 | - | 1.2 | 1.201 (0.002) |
| | G-E & E-E correlations | 0.1 | - | 1.2 | 1.208 (0.002) |
| **RxE without GxE** | No correlation | - | 0.1 | 1.2 | 1.200 (0.002) |
| | G-E correlation | - | 0.1 | 1.2 | 1.201 (0.002) |
| | E-E correlation | - | 0.1 | 1.2 | 1.198 (0.002) |
| | G-E & E-E correlations | - | 0.1 | 1.2 | 1.205 (0.002) |
| **Both GxE & RxE** | No correlation | 0.1 | 0.1 | 1.4 | 1.396 (0.002) |
| | G-E correlation | 0.1 | 0.1 | 1.4 | 1.406 (0.002) |
| | E-E correlation | 0.1 | 0.1 | 1.4 | 1.402 (0.002) |
| | G-E & E-E correlations | 0.1 | 0.1 | 1.4 | 1.416 (0.002) |
| **Kurtosis=6 and skewness=1 (non-normal distribution) for environmental variable** | | | | | |
| **No GxE & RxE** | No correlation | - | - | 1.0 | 1.001 (0.002) |
| | G-E correlation | - | - | 1.0 | 1.029 (0.002) |
| | E-E correlation | - | - | 1.0 | 1.000 (0.002) |
| | G-E & E-E correlations | - | - | 1.0 | 1.027 (0.002) |
| **GxE without RxE** | No correlation | 0.1 | - | 1.5 | 1.488 (0.007) |

| | | | | | |
|---|---|---|---|---|---|
| | G-E correlation | 0.1 | - | 1.5 | 1.520 (0.006) |
| | E-E correlation | 0.1 | - | 1.5 | 1.486 (0.006) |
| | G-E & E-E correlations | 0.1 | - | 1.5 | 1.524 (0.007) |
| **RxE without GxE** | No correlation | - | 0.1 | 1.5 | 1.500 (0.007) |
| | G-E correlation | - | 0.1 | 1.5 | 1.531 (0.008) |
| | E-E correlation | - | 0.1 | 1.5 | 1.497 (0.008) |
| | G-E & E-E correlations | - | 0.1 | 1.5 | 1.527 (0.007) |
| **Both GxE & RxE** | No correlation | 0.1 | 0.1 | 2.0 | 1.938 (0.010) |
| | G-E correlation | 0.1 | 0.1 | 2.0 | 2.003 (0.012) |
| | E-E correlation | 0.1 | 0.1 | 2.0 | 1.980 (0.011) |
| | G-E & E-E correlations | 0.1 | 0.1 | 2.0 | 2.019 (0.014) |
| **Kurtosis= 8.11 and skewness= 2.66 for binary environmental variable (k=0.1)** | | | | | |
| **No GxE & RxE** | No correlation | - | - | 1.0 | 0.999 (0.002) |
| | G-E correlation | - | - | 1.0 | 1.000 (0.002) |
| | E-E correlation | - | - | 1.0 | 0.994 (0.002) |
| | G-E & E-E correlations | - | - | 1.0 | 0.984 (0.002) |
| **GxE without RxE** | No correlation | 0.1 | - | 1.71 | 1.717 (0.004) |
| | G-E correlation | 0.1 | - | 1.71 | 1.708 (0.004) |
| | E-E correlation | 0.1 | - | 1.71 | 1.709 (0.004) |
| | G-E & E-E correlations | 0.1 | - | 1.71 | 1.703 (0.004) |
| **RxE without GxE** | No correlation | - | 0.1 | 1.71 | 1.674 (0.004) |
| | G-E correlation | - | 0.1 | 1.71 | 1.682 (0.002) |
| | E-E correlation | - | 0.1 | 1.71 | 1.678 (0.003) |
| | G-E & E-E correlations | - | 0.1 | 1.71 | 1.678 (0.003) |
| **Both GxE & RxE** | No correlation | 0.1 | 0.1 | 2.42 | 2.316 (0.005) |
| | G-E correlation | 0.1 | 0.1 | 2.42 | 2.507 (0.005) |
| | E-E correlation | 0.1 | 0.1 | 2.42 | 2.402 (0.005) |

| | | | | |
|---|---|---|---|---|
| G-E & E-E correlations | 0.1 | 0.1 | 2.42 | 2.412 (0.006) |

[a]The expected value from the theory, which is $1 + (k_{urtosis} - 1)h^2_{g_1} + (k_{urtosis} - 1)h^2_{\tau_1}$, and [b]The observed value in the GxEsum analysis. To verify the theory for the proposed model, i.e. GxEsum, we compared the obtained intercept values to the theoretical expectation under various scenarios. The estimate was obtained from 500 replicates for each scenario, which is simulated using quantitative traits. $\boldsymbol{h^2_{g_1}}$ and $\boldsymbol{h^2_{\tau_1}}$ indicates the phenotypic variance explained by GxE and RxE, respectively, i.e. When the phenotypic variance is 1, var(GxE) = $\boldsymbol{h^2_{g_1}}$ and var(RxE) = $\boldsymbol{h^2_{\tau_1}}$. All observed intercept values were consistent with the expectations under different scenarios. A non-normal distribution for environmental variables was simulated with residual values drawn from a gamma distribution with shape 0.5, which resulted in kurtosis = 6 and skewness = 1. A binary environmental variable was simulated using a liability threshold model with a population prevalence of k=0.1, which resulted in kurtosis = 8.11 and skewness = 2.66. With these non-normal or binary environmental variables, there was no inflation of type 1 error rate (Supplementary Table 3).

**Supplementary Table 3. Type I error rates of GxEsum when using non-normal environmental variable.**

| Scenarios | Type l Error rate |
|---|---|
| **When using non-normal environmental variables (a gamma distribution)** | |
| Var(GxE) = 0, Var(RxE) = 0 | 0.044 |
| Var(GxE) = 0, Var(RxE) = 0, G-E correlation = 0.1 | 0.068 |
| Var(GxE) = 0, Var(RxE) = 0, R-E correlation = 0.1 | 0.062 |
| Var(GxE) = 0, Var(RxE) = 0, G-E correlation=0.1, R-E correlation=0.1 | 0.050 |
| Var(GxE) = 0, Var(RxE) = 0.1 | 0.026 |
| Var(GxE) = 0, Var(RxE) = 0.1, G-E correlation = 0.1 | 0.036 |
| Var(GxE) = 0, Var(RxE) = 0.1, R-E correlation=0.1 | 0.050 |
| Var(GxE) = 0, Var(RxE) = 0.1, G-E correlation=0.1, R-E correlation=0.1 | 0.050 |
| **Average** | **0.048** |
| **When using binary environmental variables (k=0.1)** | |
| Var(GxE) = 0, Var(RxE) = 0 | 0.048 |
| Var(GxE) = 0, Var(RxE) = 0, G-E correlation = 0.1 | 0.046 |
| Var(GxE) = 0, Var(RxE) = 0, R-E correlation = 0.1 | 0.040 |
| Var(GxE) = 0, Var(RxE) = 0, G-E correlation=0.1, R-E correlation=0.1 | 0.056 |
| Var(GxE) = 0, Var(RxE) = 0.1 | 0.024 |
| Var(GxE) = 0, Var(RxE) = 0.1, G-E correlation = 0.1 | 0.044 |
| Var(GxE) = 0, Var(RxE) = 0.1, R-E correlation=0.1 | 0.046 |
| Var(GxE) = 0, Var(RxE) = 0.1, G-E correlation=0.1, R-E correlation=0.1 | 0.036 |
| **Average** | **0.043** |

A binary environmental variable was simulated using a liability threshold model with a population prevalence k=0.1, which resulted in kurtosis = 8.11 and skewness = 2.66. With these binary environmental, the type I error rates were quantified by repeating 500 times for each scenario.

**Supplementary Table 4. Type I error rates when using binary disease traits (k=0.1) with various confounders.**

| Scenarios | Type I error rate |
|---|---|
| Var(GxE[a]) = 0, Var(RxE[b]) = 0, G-E correlation[c]=0.1 | 0.046 |
| Var(GxE)=0, Var (RxE)=0, R-E correlation[d]=0.1 | 0.051 |
| Var(GxE)=0, Var(RxE)=0, G-E correlation=0.1, R-E correlation=0.1 | 0.046 |
| Var(GxE)=0, Var (RxE)=0.1, G-E correlation=0.1 | 0.036 |
| Var(GxE)=0, Var (RxE)=0.1, R-E correlation=0.1 | 0.044 |
| Var(GxE)=0, Var (RxE)=0.1, G-E correlation=0.1, R-E correlation=0.1 | 0.042 |
| **Average** | **0.046** |

[a]GxE: Genotype-Environment interaction, [b]RxE: Residual-Environment interaction, [c]G-E correlation: Genotype-Environment correlation, [d]R-E correlation: Residual-Environment correlation. These simulations were conducted under the null ($\sigma_{g_1}^2$=0 and $\sigma_{\tau_1}^2$=0) and RxE only ($\sigma_{g_1}^2$=0 and $\sigma_{\tau_1}^2$=0.1) scenarios with potential confounders such as G-E and/or R-E correlations. The phenotypes were standardised such that the phenotypic variance was 1. The number of replicates for each scenario was 500. The type I error rates were obtained at a significance threshold of p-value = 0.05.

**Supplementary Table 5. Type I error rates of GxEsum when the collider bias was introduced in simulated quantitative traits.**

| Scenarios | Type 1 Error Rate |
|---|---|
| Var(GxE)=0, Var(RxE)=0 | 0.062 |
| Var(GxE)=0, Var(RxE)=0, G-E correlation=0.1 | 0.064 |
| Var(GxE)=0, Var(RxE)=0, R-E correlation=0.1 | 0.04 |
| Var(GxE)=0, Var(RxE)=0, G-E correlation=0.1, R-E correlation=0.1 | 0.054 |
| Var(GxE)=0, Var(RxE)=0.1 | 0.040 |
| Var(GxE)=0, Var(RxE)=0.1, G-E correlation= 0.1 | 0.050 |
| Var(GxE)=0, Var(RxE)=0.1, R-E correlation= 0.1 | 0.066 |
| Var(GxE)=0, Var(RxE)=0.1, G-E correlation= 0.1, R-E correlation=0.1 | 0.056 |
| **Average** | **0.054** |

We further simulated by considering the potential collider bias which has been an issue in the real data analysis. The selection odds ratio was set as 1.5 and 2 for the main trait and environment, respectively (see Supplementary Note 2). After selections with collider bias, the number of samples was around 3,620 individuals, which is half of the total sample sizes. The type I error rate for each scenario was obtained from 500 replicates.

**Supplementary Table 6. Type I error rates of GxEsum in the presence of collider bias when using binary disease trait (k=0.1)**

| Scenarios | Type I error rate |
|---|---|
| Var(GxE)=0, Var(RxE)=0 | 0.058 |
| Var(GxE)=0, Var(RxE)=0, G-E correlation=0.1 | 0.050 |
| Var(GxE)=0, Var(RxE)=0, R-E correlation=0.1 | 0.038 |
| Var(GxE)=0, Var(RxE)=0, G-E correlation=0.1, R-E correlation=0.1 | 0.040 |
| Var(GxE)=0, Var(RxE)=0.1 | 0.054 |
| Var(GxE)=0, Var(RxE)=0.1, G-E correlation= 0.1 | 0.052 |
| Var(GxE)=0, Var(RxE)=0.1, R-E correlation= 0.1 | 0.044 |
| Var(GxE)=0, Var(RxE)=0.1, G-E correlation= 0.1, R-E correlation=0.1 | 0.044 |
| **Average** | **0.048** |

We further simulated binary phenotypes in the presence of collider bias (see Supplementary Note 2). After the selection based on the collider model (Supplementary Note 2), the number of samples was around 3,620 individuals, which is half of the total sample sizes. The type I error rate for each scenario was obtained from 500 simulation replicates.

**Supplementary Table 7. Comparison of the standard error (SE) of estimated GxE variance, obtained from the GCTA-GREML power calculator and from the information matrix in the RNM.**

| Sample Size | Theoretical SE[a] | Estimated SE from RNM[b] |
|---|---|---|
| **5,000** | 0.063 | 0.068 |
| **10,000** | 0.032 | 0.034 |
| **25,000** | 0.013 | 0.013 |
| **50,000** | 0.006 | 0.006 |

[a]Theoretical SE obtained from GCTA-GREML power calculator [5].
[b]Estimated SE from RNM using MTG2 software [6].

Theoretical and estiamted SE are agreed well, indicating that the theoretical SE is a good approximation of the SE of GxE varaince estiamte from RNM.

**Supplementary Table 8. Comparison of computing time between RNM and GxEsum approaches in the GxE estimation.**

| Sample size | GRM calculation | | URNM | | GWAS | | GxEsum | |
|---|---|---|---|---|---|---|---|---|
| | Time | RAM[a] | Time | RAM | Time | RAM | Time | RAM |
| **5,000** | 27 min | 0.16 GB | 6 min | 1.92 GB | 1 min | 0.27 GB | < 1 min | < 1 GB |
| **10,000** | 97 min | 0.58 GB | 30 min | 7.55 GB | 1 min | 0.42 GB | < 1 min | < 1 GB |
| **25,000** | 615 min | 3.52 GB | 227 min | 46.80 GB | 5 min | 0.86 GB | < 1 min | < 1 GB |
| **50,000** | 2537 min | 14.00 GB | 1742 min | 186.70 GB | 10 min | 1.65 GB | < 1 min | < 1 GB |
| **100,000** | NA[b] | NA | NA | NA | 21 min | 3.17 GB | < 1 min | < 1 GB |
| **200,000** | NA | NA | NA | NA | 39 min | 6.23 GB | < 1 min | < 1 GB |
| **288,837** | NA | NA | NA | NA | 66 min | 8.95 GB | < 1 min | < 1 GB |

[a]Peak random access memory (RAM) usage per each process
[b]Computing time was not recorded because it was unreasonably taken long.

Those computing requirements (time and memory) were estimated using a single CPU with a base clock speed of 2.6 GHz in sequential computing. The GxEsum and RNM analyses were conducted by using LDSC and MTG2 software, respectively.

**Supplementary Table 9. Estimates obtained from LDSC and GxEsum analyses using real data**

| Main trait | Environmental variable | Main additive genetic variance from LDSC | Main additive genetic variance from GxEsum |
|---|---|---|---|
| **BMI**[a] | Age | 0.216 (0.007) | 0.216 (0.007) |
| | NEU[b] | 0.216 (0.007) | 0.216 (0.007) |
| | PA[c] | 0.218 (0.007) | 0.218 (0.007) |
| | ALC[d] | 0.216 (0.007) | 0.216 (0.007) |
| **Hypertension** | BMI | 0.152 (0.008) | 0.152 (0.008) |
| | WHR[e] | 0.154 (0.008) | 0.154 (0.008) |
| | BFP[f] | 0.151 (0.008) | 0.151 (0.008) |
| **Type 2 Diabetes** | BMI | 0.142 (0.014) | 0.141 (0.014) |
| | SBP[g] | 0.198 (0.014) | 0.198 (0.014) |
| | DBP[h] | 0.204 (0.014) | 0.204 (0.014) |

[a]Body Mass Index
[b]Neuroticism
[c]Physical activity
[d]Alcohol Intake frequency
[e]Waist Hip ratio
[f]Body fat percentage
[g]Systolic blood pressure
[h]Diastolic blood pressure

We used a quantitative trait (BMI) and a binary disease trait (hypertension and type 2 diabetes) because BMI is known to be modulated by age/lifestyle such as NEU, ALC, PA [7-9], and hypertension and type 2 diabetes are known to be caused by obese traits such as BMI and WHR[10,11]. The p-value is from a Wald test for the estimated GxE variance not being different from zero. The estimates on the observed scale for the binary trait, hypertension, were transformed to those on the liability scale using Robertson transformation[12,13]. The column for LDSC is obtained main genetic variances using LDSC approach (see Supplementary Note 4 for the model description).

**Supplementary Table 10. Obtained p-values for GxE in BMI with 4 covariates with and without phenotypic imputation**

| | | Conditions | N[a] | P-value[b] |
|---|---|---|---|---|
| NEU | Model 1 | BMI adjusted for demographics[c], 10 PCs[d], age and NEU[e] | 231,889 | 3.43E-05 |
| | Model 2 | BMI adjusted for demographics, 10 PCs, age and NEU with imputed phenotypic data | 288,837 | 1.61E-05 |
| ALC[f] | Model 1 | BMI adjusted for demographics, 10 PCs, age and ALC | 205,505 | 2.25E-01 |
| | Model 2 | BMI adjusted for demographics, 10 PCs, age and ALC with imputed phenotypic data | 288,837 | 5.98E-02 |
| Age | Model 1 | BMI adjusted for demographics 10 PCs and age | 284,340 | 1.59E-02 |
| | Model 2 | BMI adjusted for demographics 10 PCs and age with imputed phenotypic data | 288,837 | 1.86E-02 |
| PA[g] | Model 1 | BMI adjusted for demographics, 10 PCs, age and PA | 231,972 | 1.06E-02 |
| | Model 2 | BMI adjusted for demographics, 10 PCs, age and PA with imputed phenotypic data | 288,837 | 2.57E-02 |

[a]The number of sample sizes used in the analysis
[b]Estimated P-value for GxE
[c]Demographics include sex, centre, Townsend Deprivation Index, genotype batch, year of birth, income and education.
[d]First 10 principle components
[e]Neuroticism score
[f]Alcohol intake frequency
[g]Physical activity

**Model 1:** The first model is using the phenotypes adjusted for demographics (sex, centre, Townsend deprivation index, batch, year of birth, income and education), 10 PCs and the environmental variable that is used in the analysis
**Model 2:** The second model is the same as the model 1 except that missing phenotypes are imputed with the mean value for each variable so that the number of samples in model 2 is larger than model 1.

Note that the missing rate for the main phenotypes (BMI) is 0.0032.

**Supplementary Table 11. Obtained p-values for GxE in Hypertension with 3 covariates with and without phenotypic imputation**

| | | Conditions | N | P-value |
|---|---|---|---|---|
| BMI | Model 1 | Hypertension adjusted for demographics, 10 PCs, age and BMI | 262,220 | 1.60E-03 |
| | Model 2 | Hypertension adjusted for demographics, 10 PCs, age and BMI using imputed phenotypic data | 288,837 | 2.09E-03 |
| BFP[a] | Model 1 | Hypertension adjusted for demographics, 10 PCs, age and BFP | 258,424 | 8.24E-02 |
| | Model 2 | Hypertension adjusted for demographics, 10 PCs, age and BFP using imputed phenotypic data | 288,837 | 2.66E-02 |
| WHR[b] | Model 1 | Hypertension adjusted for demographics, 10 PCs, age and WHR | 262,556 | 4.55E-02 |
| | Model 2 | Hypertension adjusted for demographics, 10 PCs, age and WHR using imputed phenotypic data | 288,837 | 3.21E-02 |

[a]Body fat percentage
[b]Waist-hip ratio was obtained from waist circumference / hip circumference

**Model 1:** The first model is using the phenotypes adjusted for demographics (sex, centre, Townsend deprivation index, batch, year of birth, income and education), 10 PCs and the environmental variable that is used in the analysis
**Model 2:** The second model is the same as the model 1 except that missing phenotypes are imputed with the mean value for each variable so that the number of samples in model 2 is larger than model 1.

Note that the missing rate for the main phenotypes (hypertension) is 0.079.

**Supplementary Table 12. Obtained p-values for GxE in Type 2 diabetes with 3 covariates with and without phenotypic imputation**

| | | Conditions | N | P-value |
|---|---|---|---|---|
| BMI | Model 1 | Type 2 diabetes adjusted for demographics, 10 PCs, age and BMI | 283,784 | 2.01E-04 |
| | Model 2 | Type 2 diabetes adjusted for demographics, 10 PCs, age and BMI using imputed phenotypic data | 288,837 | 1.58E-04 |
| SBP[a] | Model 1 | Type 2 diabetes adjusted for demographics, 10 PCs, age and SBP | 259,700 | 4.63E-01 |
| | Model 2 | Type 2 diabetes adjusted for demographics, 10 PCs, age and SBP using imputed phenotypic data | 288,837 | 5.38E-01 |
| DBP[b] | Model 1 | Type 2 diabetes adjusted for demographics, 10 PCs, age and DBP | 259,705 | 3.51E-01 |
| | Model 2 | Type 2 diabetes adjusted for demographics, 10 PCs, age and DBP using imputed phenotypic data | 288,837 | 3.17E-01 |

[a]Systolic blood pressure
[b]Diastolic blood pressure

**Model 1:** The first model is using the phenotypes adjusted for demographics (sex, centre, Townsend deprivation index, batch, year of birth, income and education), 10 PCs and the environmental variable that is used in the analysis
**Model 2:** The second model is the same as the model 1 except that missing phenotypes are imputed with the mean value for each variable so that the number of samples in model 2 is larger than model 1.

Note that the missing rate for the main phenotypes (type 2 diabetes) is 0.002.
